# A Spatiotemporal Map of Co-Receptor Signaling Networks Underlying B Cell Activation

**DOI:** 10.1101/2023.03.17.533227

**Authors:** Katherine J. Susa, Gary A. Bradshaw, Robyn J. Eisert, Charlotte M. Schilling, Marian Kalocsay, Stephen C. Blacklow, Andrew C. Kruse

## Abstract

The B cell receptor (BCR) signals together with a multi-component co-receptor complex to initiate B cell activation in response to antigen binding. This process underlies nearly every aspect of proper B cell function. Here, we take advantage of peroxidase-catalyzed proximity labeling combined with quantitative mass spectrometry to track B cell co-receptor signaling dynamics from 10 seconds to 2 hours after BCR stimulation. This approach enables tracking of 2,814 proximity-labeled proteins and 1,394 quantified phosphosites and provides an unbiased and quantitative molecular map of proteins recruited to the vicinity of CD19, the key signaling subunit of the co-receptor complex. We detail the recruitment kinetics of essential signaling effectors to CD19 following activation, and then identify new mediators of B cell activation. In particular, we show that the glutamate transporter SLC1A1 is responsible for mediating rapid metabolic reprogramming immediately downstream of BCR stimulation and for maintaining redox homeostasis during B cell activation. This study provides a comprehensive map of the BCR signaling pathway and a rich resource for uncovering the complex signaling networks that regulate B cell activation.

## Introduction

B cells are central components of the adaptive immune system. They respond to infectious agents by producing and secreting antibodies, yet dysregulation of the B cell response can result in autoimmune disease or B cell malignancies^1^. Antibodies produced by B cells are also important therapeutics and tools in biomedical research. In addition to secreting antibodies, B cells produce immunomodulatory cytokines, guide the organization and development of lymphoid structures, and provide co-stimulatory signals to T cells^1^. Central to this multifaceted biology is the transition of the B cell from a quiescent or “resting” state to an activated state in response to an antigen. Activation is initiated when an antigen binds to a cell surface B cell receptor (BCR), which leads to a multi-branched signaling cascade that culminates in changes in cell metabolism, cytoskeletal reorganization, the upregulation of co-stimulatory, migration, and adhesion receptors, cell-cycle progression, and ultimately, differentiation into an antibody-secreting effector cell^2, 3^.

Not surprisingly, aberrant BCR signaling is closely associated with B cell dysfunction. For example, excessive BCR signaling is a hallmark of B cell-driven autoimmune diseases like rheumatoid arthritis or systemic lupus erythematosus^4, 5^. Chronic BCR signaling is also associated with various B cell-derived malignancies^6, 7^. Accordingly, B cell surface proteins, including CD20 and CD19, and downstream kinases within the BCR signaling pathway have become important targets for therapeutics that have transformed the treatment of B cell cancers^8, 9^ and B cell-driven autoimmune disorders^10, 11^.

The precise regulation and molecular coordination of signaling effectors during B cell activation underlies proper B cell function. Current methods for tracking BCR signaling have illuminated parts of the complex, multi-branched, and dynamic signaling events underlying activation, but have been limited by poor temporal resolution, low throughput, and the frequently required use of overexpression for candidate-based approaches. While previously reported co-immunoprecipitation, knockout, and live cell microscopy experiments^12–14^ have identified major intracellular signaling pathways underlying B cell activation, many aspects of BCR signaling remain incompletely characterized. Most notably, very little is known about the dynamics and relevant proteins involved in the earliest signaling events that occur in the seconds and minutes following antigen binding to the BCR.

CD19 is the essential signaling subunit of a multi-component co-receptor complex that functions in concert with the BCR to execute the cellular response to antigen. This complex, which also includes the complement receptor CR2, the tetraspanin CD81, and Leu-13 (also known as IFITM1), enhances the sensitivity of B cell signaling by up to 1000-fold to enable B cells to respond to low concentrations of antigen ^15, 16^. Upon BCR stimulation, the large cytoplasmic tail of CD19 undergoes phosphorylation at highly conserved tyrosine residues on the cytoplasmic tail, creating docking sites for the Src homology 2 (SH2) domains of important meditators of BCR signaling^17^, including phosphatidylinositol 3-kinase (PI3K), Vav, phospholipase C (PLC), and protein tyrosine kinases such as Lyn^18–22^. Thus, CD19 serves as a membrane-anchored adaptor protein that modulates the response of several activation-dependent signals, leading to sustained signaling of pathways necessary for B cell activation.

The phenotypes of CD19 mutations also highlight the necessity for CD19 signaling in B cell activation. CD19-deficient mice are unresponsive to antigen receptor stimulation^23^, and humans with CD19 mutations are severely immunodeficient^23, 24^. In contrast, mice over-expressing CD19 show striking defects in early B cell development, augmented response to stimulation, and autoimmune-like phenotypes, such as spontaneous secretion of auto-antibodies^23, 25^.

Here, we took advantage of the importance of CD19 as a critical node in BCR signaling to map the response to a BCR stimulus in an unbiased and quantitative manner using multiplexed proximity proteomics **(Figure 1)**. We expressed CD19 fused to the engineered ascorbate peroxidase APEX2^26, 27^ from its endogenous locus in Raji B cells **(Figure S1)**, and analyzed the pattern of biotinylated CD19-proximal proteins as a function of time to produce a dynamic view of the cellular response to BCR stimulation. Using this approach, we were able to define the kinetics of events implicated in the signaling response to BCR engagement, including recruitment- and phosphorylation-kinetics of protein kinase C (PKC) isoforms, PI3K, Akt, and isoforms of Raf. We identified other proteins responsive to BCR stimulation that have not been characterized previously, including the glutamate transporter SLC1A1. We found that SLC1A1 mediates rapid metabolic preprogramming during the B cell response, highlighting the valuable resource our spatio-temporally resolved dataset provides for uncovering new B cell biology and for identifying potential targets to modulate B cell activation. Precise kinetics and measurement of local, in contrast to traditionally measured global, phosphorylation events display molecular events in CD19 signaling *in vivo* and *in situ* at unprecedented resolution, allowing for direct mapping of signaling effectors recruited to CD19 throughout BCR signaling.

**Figure 1.**
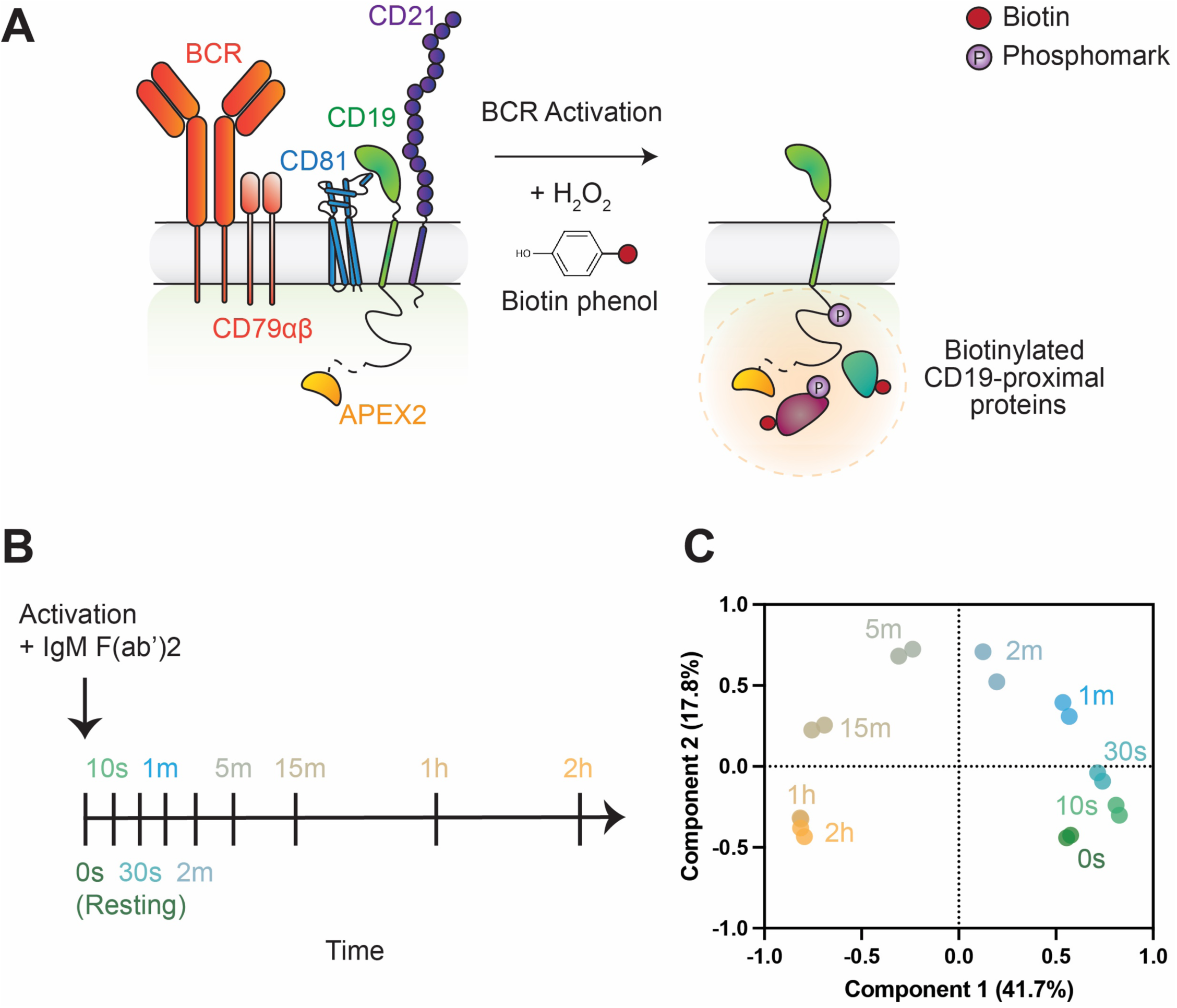
Overview of CD19-APEX system. **(A)** Schematic of CD19-APEX proximity labeling. CD19, the signaling subunit of the B cell co-receptor complex, is tagged with the ascorbate peroxide APEX2 on its C- terminus. In the presence of hydrogen peroxide (H_2_O_2_), APEX2 converts biotin phenol into a biotin phenoxy radical, which then covalently labels proteins within close spatial proximity (about 20 nm) of the APEX2 fusion. **(B)** Timeline of the eight time points at which CD19-APEX cells were labeled after activation. **(C)** Principal component analysis of biological replicates.

## Results

### Engineering and Validation of CD19-APEX2 B Cells

To achieve inducible *in vivo* labeling of proteins within a small radius of 10-20 nm around CD19, we created a CD19-APEX2 fusion protein, in which APEX2 is fused to CD19 at its carboxy terminus (C-terminus) using a flexible glycine-serine (GS) repeat linker **(Figure 1A; Figure S1A)**. Importantly, the CD19 C-terminus is predicted to be primarily unstructured^28^, suggesting that an APEX2 fusion would be unlikely to alter function.

Properly folded CD19 is trafficked to the cell surface by forming a complex with the tetraspanin CD81^29^. As an initial test for the functional integrity of our fusion construct, we used a trafficking assay in HEK293T cells to assess the ability of CD81 to enhance the delivery of CD19-APEX2 to the cell surface^30, 31^. Our CD19-APEX2 protein matched the response of wild-type CD19, showing a three-fold increase in cell surface staining when co-transfected with CD81 **(Figure S1B, C)**, confirming that the fusion protein was properly transported to the cell surface and that its interaction with CD81 responsible for surface export was intact.

In previous reports of proximity labeling experiments using APEX2, stable cell lines were generated in which APEX2 fusions were overexpressed. Examples of such studies include reports that tracked GPCR signaling^32, 33^, mapped the proteome of the mitochondrial matrix^34^ and cilia^35^, or monitored signaling from lipid rafts in B cells^36^. However, a similar approach in which the B cell co-receptor is ectopically overexpressed would be problematic because murine B cells overexpressing CD19 are hyper-responsive to BCR stimulation, resulting in abnormal B cell signaling, suppression of peripheral tolerance and secretion of autoantibodies^37^. Instead, here we used CRISPR/Cas9 genome editing to insert APEX2 at the C terminus of CD19 at the endogenous locus in Raji B cells. This widely used B cell line expresses the IgM BCR isotype on its cell surface^38^. Our engineered cells are homozygous for the CD19-APEX2 knock-in **(Figure S1D),** and this knock-in has no detectable off-target integrations in the genome **(Figure S1E)**. While the fusion shows slightly reduced surface staining on the B cell surface compared to parental Raji cells, the amount of CD19 expressed is within the range expressed on primary human B cells and parental Raji cells, indicating that CD19 is present in physiologically relevant amounts in our engineered cells **(Figure S1F)**.

Next, we confirmed that our CD19-APEX2 cell line responds normally to BCR stimulation. Within hours to days after activation, B cells upregulate the surface expression of co-stimulatory receptors, including the canonical activation markers CD69 and CD86^39, 40^. After BCR stimulation with an anti-IgM crosslinking Fab fragment (α-IgM), an activating reagent that crosslinks the BCR to mimic T cell-independent activation by oligomeric antigens, our engineered cells showed a three-fold increase in CD69 surface staining and two-fold increase CD86 surface staining, matching the profile of parental Raji cells **(Figure S2A)**. We also assessed whether this functional signaling response depended on the presence of CD19. We created CD19 knockout Raji cells (**Figure S2B, C**) and monitored the ability of these cells to induce elevated amounts of CD69 and CD86 on their cell surface upon BCR stimulation with α-IgM **(Figure S2A)**. All three CD19 knockout clones failed to increase the surface expression of these markers, indicating that our CD19-APEX2 fusion preserves functional BCR signaling, whereas CD19 knockout eliminates this downstream response in Raji cells. To assess the functional integrity of proximal, early signaling events, we monitored the induction of phosphorylation on Tyrosine 531 (Y531) of the cytoplasmic tail of CD19 upon activation. This essential regulatory mark bridges CD19 to phosphoinositide 3-kinase (PI3K) and Src family tyrosine kinases to enable downstream activation of these pathways^19, 41, 42^. After 2 minutes of BCR crosslinking, Y531 of CD19 is phosphorylated in both parental and CD19-APEX2 cells, indicating that our CD19-APEX2 fusion is functional in initiating signaling **(Figure S2D)**.

Importantly, because the APEX2 reaction requires the addition of peroxide to cells, and peroxide has been shown to trigger B cell signaling through inhibition of protein phosphatases^43^, we monitored phosphorylation upon the addition of peroxide to Raji cells using an antibody that detects a broad range of phosphorylated proteins. In line with these previous reports, we observed a significant increase in phosphorylation upon the addition of 10 mM peroxide but observed no increase in phosphorylation with the amount of peroxide used in our APEX2 experiments (0.25 mM), indicating that signaling is not triggered by the labeling reaction alone **(Figure S2E)**.

### Time-resolved signaling from 10 seconds to 2 hours after BCR Stimulation

Having validated that our CD19-APEX2 cell line exhibits normal hallmarks of BCR signaling, we then sought to assess the CD19 microenvironment after BCR stimulation. The B cell signaling response is rapid and dynamic, forming short-lived signaling complexes that initiate downstream signaling through processive phosphorylation and ubiquitylation of signaling partners. Previous studies of early events in this response were limited by poor temporal resolution and the difficulty to monitor transient signaling events. These technical obstacles precluded a complete understanding of the identity, timing, and order of signaling events during the activation response^21, 44, 45^. Using our CD19-APEX2 system (**Figure 1A)**, we tracked CD19 interactions at eight timepoints after BCR stimulation, ranging from 10 seconds to 2 hours after the addition of an α-IgM F(ab’)2 fragment **(Figure 1B; Figure S3A)**. After streptavidin affinity purification and analysis of resulting peptides by triple-stage mass spectrometry (MS^3^), this time-resolved analysis allowed tracking of CD19 proximity kinetics for 2,814 proteins and 1,394 phosphosites (**Figure S3B**). For data analysis, protein and phosphosite measurements were combined and normalized to the enrichment of CD19 at each timepoint. This experiment yielded a robust data set, with measurements of protein abundance strongly correlated (R^2^>0.99) across each pair of biological replicates, indicating highly reproducible protein quantification **(Figure S4).** Additionally, principal component analysis shows that the replicates group tightly together with clear separation among time points, and with samples showing a sequential movement in principal component space during the activation time course **(Figure 1C)**. In an accompanying experiment, we also included reference samples prepared without adding biotin phenol or peroxide alongside a sparser time course of labeling, allowing us to assess the extent of endogenous biotinylation and non-specific labeling in the samples. **(Figure S5A)**. By comparing the abundance of peptides derived from CD19-APEX2 cells treated with peroxide and biotin phenol to those without treatment, we confirmed that known CD19 interactors were highly enriched in our data set by comparison to endogenously biotinylated proteins **(Figure S5B)**. The core components of the co-receptor complex, CR2 and CD81, were among the most highly enriched proteins. Other key signaling effectors, such as Syk and Lyn kinases, also showed strong enrichment. Overall, over 96% of all identified proteins showed at least a five-fold enrichment over no biotin phenol and peroxide controls, indicating highly specific biotinylation restricted to the CD19 microenvironment **(Figure S5C).** Because of this highly specific labeling pattern, likely because we inserted the APEX2 fusion into the endogenous CD19 locus to ensure physiologic expression levels, we omitted the no biotin phenol and peroxide samples in the complete 18-plex time course in order to capture a more comprehensive sampling of early time points after BCR stimulation **(Figure 1B)**.

One limitation of our approach is that a protein proximal to the APEX2 fusion may not be detected for one of several technical reasons, including steric obstruction of electron-rich residues (e.g., tyrosine), a paucity of reactive residues in the protein sequence, or inherent unsuitability of the protein for analysis by mass spectrometry. For example, we did not detect IFITM1 (also known as Leu-13), an interferon-inducible membrane protein reported to interact with CR2, CD19, and CD81^15, 46^. A possible reason for MS analysis having missed IFITM1 is that tryptic digest of this small protein with only 125 amino acids produces few peptide species suited for MS analysis.

Our dataset shows both the kinetics of recruitment of proteins previously described as part of the BCR signaling pathway and those without a previously defined role in B cell activation. We established a micro-enrichment workflow for phosphopeptides from the same samples, allowing us to quantify several hundred phosphosites in addition to relative protein levels from the same TMT multiplexes. Proteins or phosphopeptides with a fold change difference of at least 2.5 in stimulated cells compared to resting cells show at least five distinct response patterns by hierarchical clustering **(Figure 2A)**, with representative examples from each response pattern group shown in Figures 2B-F. Among the proteins or phosphosites exhibiting enrichment in labeling upon activation, increased labeling can be observed at early (<5 min.) time points (**Figure 2B**), late (>15 min.) time points **(Figure 2C)**, or throughout the time course **(Figure 2D)**. On the other hand, other proteins or phosphosites showed reduced labeling at different times after activation, with early depletion of proteins like the serine/threonine kinase MARK2 and its quantified phosphosite on S456 following the same trend **(Figure 2E)**, or late depletion like the thioredoxin-interacting protein TXNIP and its phosphosite S308 **(Figure 2F)**.

**Figure 2.**
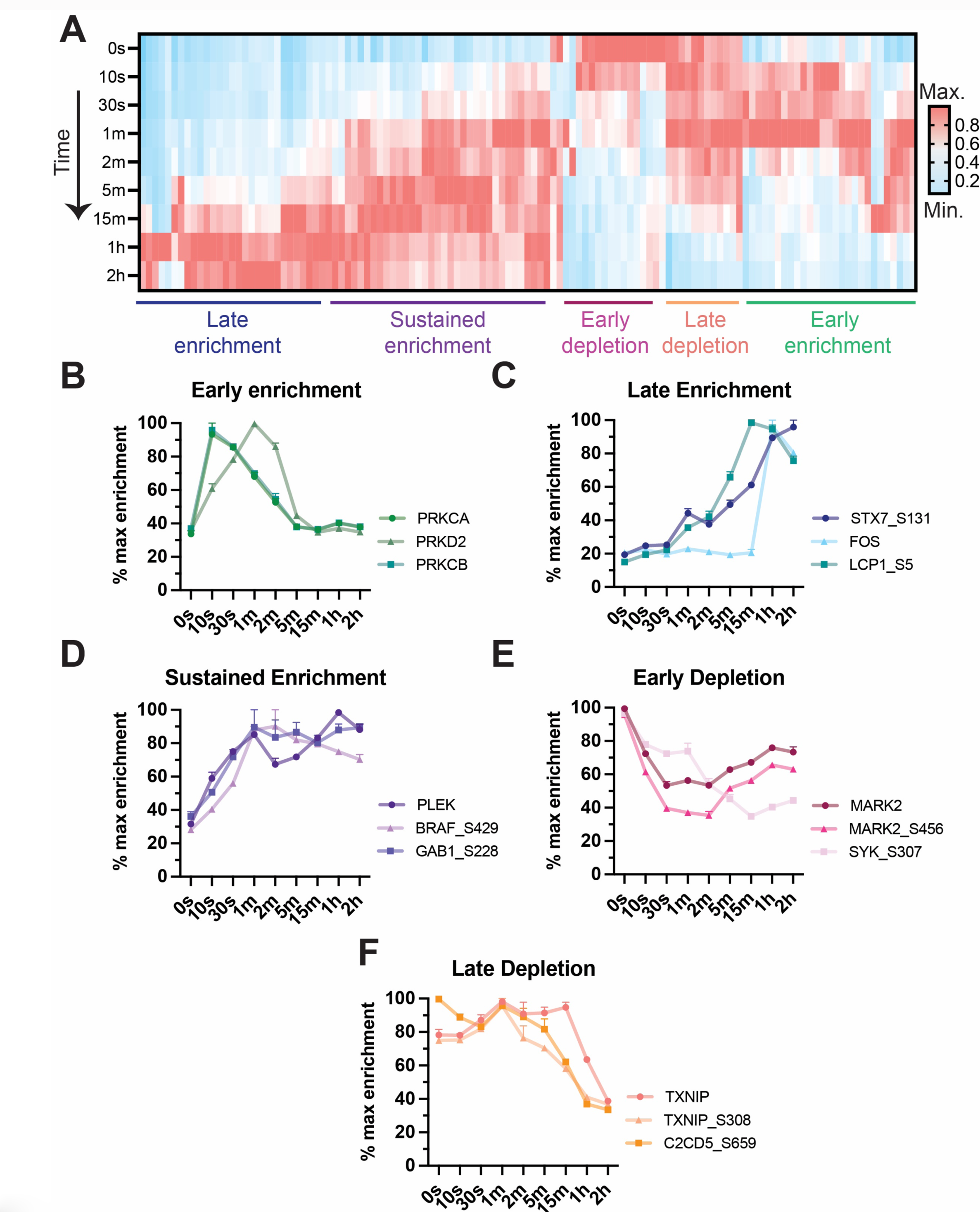
Response patterns to BCR stimulation. **(A)** Enrichment pattern heat map of all proteins and phosphosites with a fold change difference of at least 2.5 upon activation, highlighting five distinct patterns of response revealed by hierarchical clustering. Each protein’s enrichment level across different time points is normalized to its maximum enrichment signal. Enrichment profile of three exemplary proteins showing: **(B)** an early enrichment response pattern, **(C)** a late enrichment response pattern, **(D)** a sustained enrichment response pattern, **(E)** an early depletion response pattern, and **(F)** a late depletion response pattern. Error bars represent mean ± SEM of two biological replicates. In all panels, phospho-marks are annotated with the protein name followed by “_amino acid position” of the phosho-mark.

### Kinetics of BCR signaling core components to CD19

Our time-resolved CD19-APEX2 proximity labeling data allowed us to determine the order and timing of recruitment of previously known essential signaling effectors to CD19 upon activation **(Figure 3A and Figure 3B)**. While others previously showed that these signaling effectors are involved in activation, our study revealed dynamic engagement of different proteins with CD19, defining the kinetics of transient, CD19-dependent signal transduction events. We detected a rapid and temporary increase in the recruitment of members of the protein kinase C family, phosphorylation of Syk and BLNK, and recruitment of essential second messenger generators beginning as early as 10 seconds after BCR stimulation **(Figure 3C and 3D)**. At later time points, we observed sustained recruitment of effectors in the Akt and Raf-MAPK pathways, along with the inhibitory co-receptor CD22 **(Figure 3C and 3D).**

**Figure 3.**
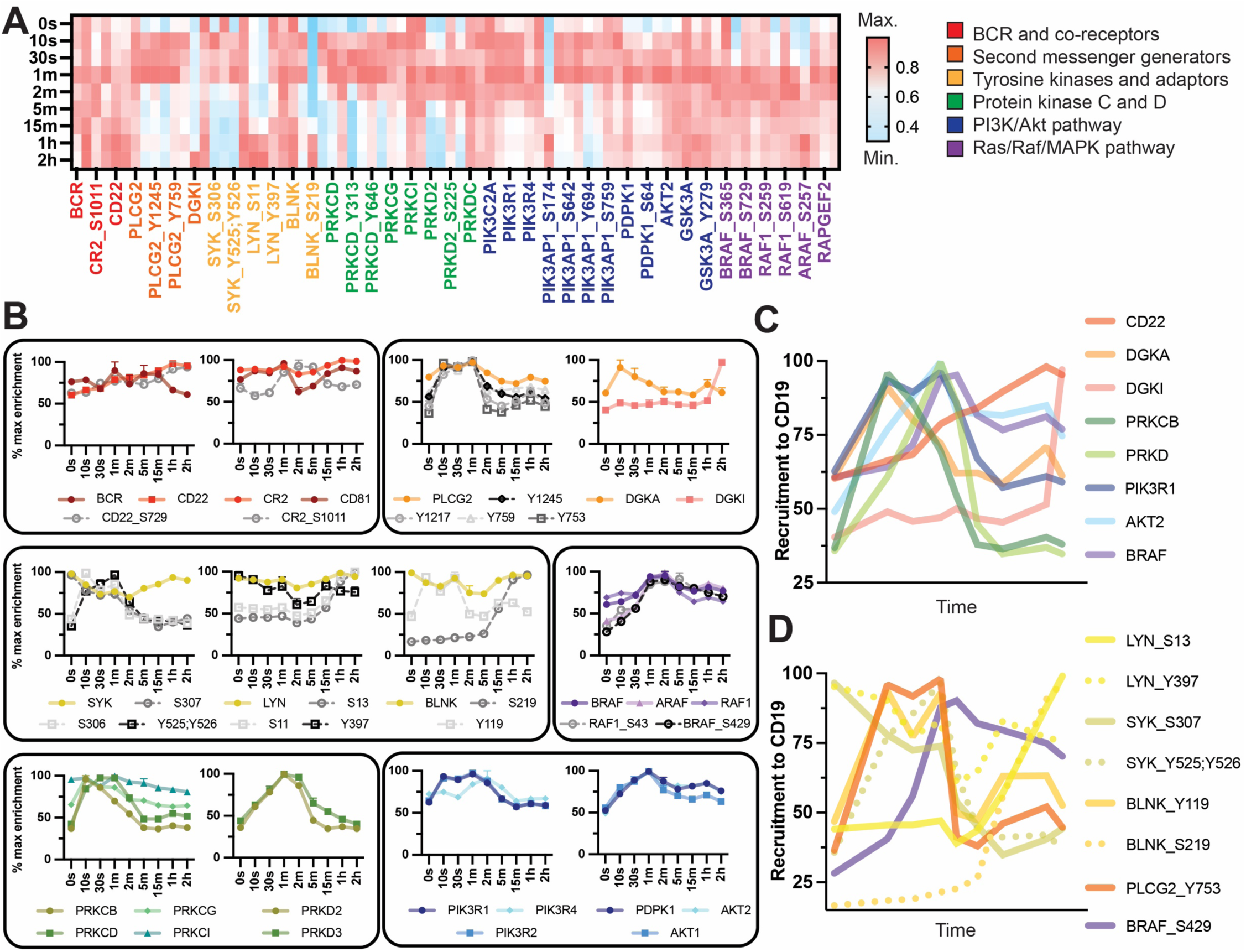
Kinetics of recruitment of core components of the BCR signaling pathway to CD19. **(A)** Enrichment pattern heat map of known components of the BCR signaling pathway, grouped by pathway. **(B)** Enrichment profiles of known components of the BCR signaling pathway, highlighting the distinct patterns of response of each member. Error bars represent mean ± SEM of two biological replicates, and dashed grey lines represent phosphomarks. **(C)** Summary of the proteins recruited to CD19 during activation. Time is plotted on a log10 scale for better separation of the early time points. **(D)** Summary of phosphorylation within the CD19 microenvironment during activation. Time is plotted on a log10 scale for better separation of the early time points. Panels C-D contain the same data shown in the Panel A heatmap plotted in different ways to highlight distinct patterns of response. In all panels, phospho-marks are annotated with the protein name followed by “_amino acid position” of the phosho-mark.

One of the first events we observed after BCR stimulation was robust and transient enrichment of phosphorylated Tyrosine 525/526 (Y525/526) of Syk, a tyrosine kinase that phosphorylates members of the protein kinase C (PRKC, also known as PKC) family, phospholipase C-γ2 (PLCG2), and B cell linker protein (BLNK) to drive early events in the BCR signaling pathway. While Syk total protein enrichment does not increase in proximity of CD19 upon BCR stimulation, Y525/526 of Syk are rapidly phosphorylated and remain phosphorylated from 10 seconds to 2 minutes after BCR stimulation **(Figure 3B, third panel)**. Y525/526 are located within the activation loop of the Syk kinase domain^47^, and their rapid phosphorylation suggests that while Syk is present in the CD19 microenvironment both before and after BCR stimulation, it becomes rapidly and transiently phosphorylated to facilitate its activation during the first two minutes of activation. In line with this, we observe rapid and transient phosphorylation of the Syk substrates, PLCG2 and BLNK **(Figure 3B, third panel)**. BLNK is an adaptor protein that, when phosphorylated, functions to provide a scaffold for assembling complexes involved in downstream BCR signaling, including those containing PLCG2^48^. While the enrichment of BLNK does not change substantially upon BCR stimulation, we observed a striking enrichment of Y119 phosphorylation on BLNK. This phospho-mark becomes rapidly enriched 10 seconds after activation and then returns to basal levels at 2 minutes **(Figure 3B, third panel)**. In contrast, phosphorylation of S219 on BLNK appears only at late time points, with strong enrichment 1 and 2 hours after activation, suggesting that dynamic changes in phosphorylation attenuate BLNK signaling at distinct time points during activation **(Figure 3B, third panel)**. In the same 10-second to 2-minute time interval, PLCG2 is transiently phosphorylated on tyrosine at positions 753, 759, 1217, and 1245 **(Figure 3B, second panel)**. These marks have been shown by others to increase phospholipase activity, leading to the production of the critical second messengers diacylglycerol (DAG) and inositol 1,4,5-trisphosphate (IP3)^49^. The generation of IP3 leads to enzyme activation and calcium influx from the endoplasmic reticulum, while DAG activates the protein kinase C family (PRKC, also known as PKC), which are critical drivers of BCR-induced NF-KB activation, controlling cytokine production, gene regulation, and B cell-survival^50^.

In line with the production of DAG beginning 10 seconds after stimulation, we observe rapid recruitment of several PKC isoforms to CD19, with enrichment peaking at 10 seconds and returning to basal levels after 5 minutes **(Figure 3B, fifth panel)**. At the same time as PKC isoforms are recruited to CD19, the diacylglycerol kinase DGKA is also recruited, with the enrichment profile of DGKA directly matching that of PKC, peaking at 10 seconds after activation and returning to basal levels within 5 minutes **(Figure 3B, second panel)**. DGKA catalyzes the conversion of DAG into phosphatidic acid (PA), indicating that during this time of early signal amplification, negative regulators are also recruited to CD19 to control and limit the strength of PKC activation and antigen receptor signaling. In contrast to the early enrichment of DGKA, another isoform, DGKI, is robustly recruited to CD19 beginning only at 2 hours after BCR stimulation **(Figure 3B, second panel)**. DGKI has been shown by others to promote the remodeling of the actin cytoskeleton during antigen uptake within the immune system^51^, suggesting that it is acting at later points during activation to promote functions associated with antigen processing.

Concurrent with the recruitment of PKC, DGKA, and the phosphorylation of PLGC2, PI3K is also proximity labeled by CD19 beginning at 10 seconds after BCR stimulation, and it remains proximal through 2 minutes after stimulation **(Figure 3B, sixth panel)**. This finding recapitulates early work showing that PI3K binds to a phosphotyrosine-containing motif on the CD19 cytoplasmic tail after BCR stimulation^19^. PI3K is responsible for phosphorylating phosphatidyl-inositol-4,5-triphosphate (PIP2) to generate phosphatidyl-inositol-3,4,5- triphosphate (PIP3), an essential second messenger that activates the Akt pathway, an important driver of cell cycle progression and proliferation. In line with the recruitment kinetics of PI3K, we observe robust enrichment of 3-phosphoinositide-dependent protein kinase-1 (PDPK1), the kinase responsible for phosphorylating Akt within its activation segment, beginning at 10 seconds and peaking at 1 minute after BCR stimulation **(Figure 3B, sixth panel).** The enrichment profile of PDPK1 closely matches that of its substrates, Akt1 and Akt2, suggesting that both kinases rapidly re-localize to CD19 to promote the initial activation of Akt **(Figure 3B, sixth panel)**. PDPK1, Akt1, and Akt2 enrichment begins to decrease at 2 minutes. Downstream Akt pathway proteins, such as GSK3A **(Figure 3A)**, do not show CD19-recruitment, suggesting that Akt is recruited to CD19 for its initial activation, and that then downstream effectors move to a different microenvironment, potentially to prevent unrestrained Akt pathway amplification.

Shortly after activation of the PI3K/Akt pathway, all three isoforms of Raf are rapidly recruited to the CD19 microenvironment, with enrichment beginning at 30 seconds and peaking within 2 minutes of BCR stimulation **(Figure 3B, sixth panel)**. Raf kinases play an important role in the modulation of ERK activity in the mitogen-activated protein (MAP) kinase pathway, which is used by B cells to control cellular proliferation, survival, and differentiation. While others have shown that CD19 is not essential for activating the Raf-MAPK pathway^52^, our data shows that all three Raf isoforms are still robustly recruited to the CD19 microenvironment. A potential reason for this recruitment is to provide additional regulation to allow for more precise control of Raf activation and signaling. In line with this, we observe strong enrichment of B-Raf Serine 429 (S429) phosphorylation, peaking at 2 minutes after BCR stimulation **(Figure 3B, fourth panel)**. This phospho-mark, in the N-terminal regulatory domain of B-Raf, is deposited by Akt and functions as a negative regulatory mark to decrease B-Raf activity^53^.

### CD19/BCR association dynamics with co-receptors

In addition to revealing the choreography of intracellular signaling effector recruitment to CD19, our dataset also allowed us to understand the organization of the BCR and other co-receptors in relation to CD19 during BCR stimulation. A key question in BCR signal transduction is how BCR complexes stay silent on resting cells and whether they are physically separated from activating co-receptors on the resting B cell surface. Enrichment of the IgM BCR remains relatively constant throughout the time course, decreasing in enrichment slightly at 1 and 2 hours after BCR stimulation **(Figure 3B, first panel)**. This relatively stable enrichment level aligns with previous reports using high-resolution microscopy to study the organization of the BCR, which showed that BCRs organize into nanoscale clusters on the B cell surface and that initiation of BCR signaling is not accompanied by a global alteration in BCR organization^54^.

We also monitored the dynamics of other members of the B cell co-receptor complex in response to stimulation. CD19 functions within a complex containing the complement receptor CR2 and the tetraspanin CD81. CR2 does not change enrichment throughout the time course, which is expected because we stimulated the B cells in a complement-independent manner **(Figure 3B, first panel)**. We have previously shown that CD81 dissociates from CD19 on activated B cells, measured 72 hours after activation^30^. In this study focusing on early time points, CD81 enrichment decreases at 2 minutes and then returns to basal levels, suggesting dynamic regulation throughout activation. The inhibitory co-receptor, CD22, increases in CD19-proximity throughout the time course, reaching its highest enrichment 2 hours after activation. This dynamic recruitment of different co-receptors to CD19 throughout activation indicates that the reorganization of CD19 complexes during activation facilitates amplification and then dampening of CD19-dependent signaling responses.

### Spatiotemporally resolved CD19-APEX2 data uncover new B cell biology

In addition to delineating sequence and timing of BCR signaling effector protein recruitment, we also observed striking time-dependent changes in the enrichment of proteins recruited to the B cell co-receptor not previously described in BCR signaling (**Figure 4**). First, we identified rapid induction in the enrichment of proteins involved in cytoskeletal regulation throughout the time course **(Figure 4A)**. For example, pleckstrin (PLEK) and its phospho-mark Serine117 showed robust enrichment as early as 10 seconds after BCR stimulation and remained highly enriched for the duration of the time course. Pleckstrin has no previously reported roles in B cell signaling but is known to be phosphorylated by PKC at position 117 to regulate the reorganization of the actin cytoskeleton and the formation of lamellipodia^55, 56^. Regulation of the actin cytoskeleton plays an essential role during several stages of B cell activation, including in the distribution of the BCR and co-receptors in the B cell membrane, cellular spreading, the formation of the immune synapse, and the gathering of antigens for internalization and processing^57^. Other proteins with no previously reported connection to B cell signaling include members of the microtubule affinity-regulating kinase family (MARK2 and MARK3), which are strongly depleted upon activation, and the actin-binding protein Plastin-2 (LCP1), which showed no change in total protein abundance throughout the time course, but a marked enrichment of phosphorylation of two serine residues (S5 and S7) near its N-terminus. Although there are no reports of LCP1 function in B cells, in T cells, phosphorylation of S5 is necessary for protein trafficking of the activation markers CD25 and CD69 to the cell surface^58^, and LCP1 knockout T cells are markedly deficient in cytokine secretion and proliferation^59^, consistent with the idea that LCP1 may be involved in similar functions on B cells.

**Figure 4.**
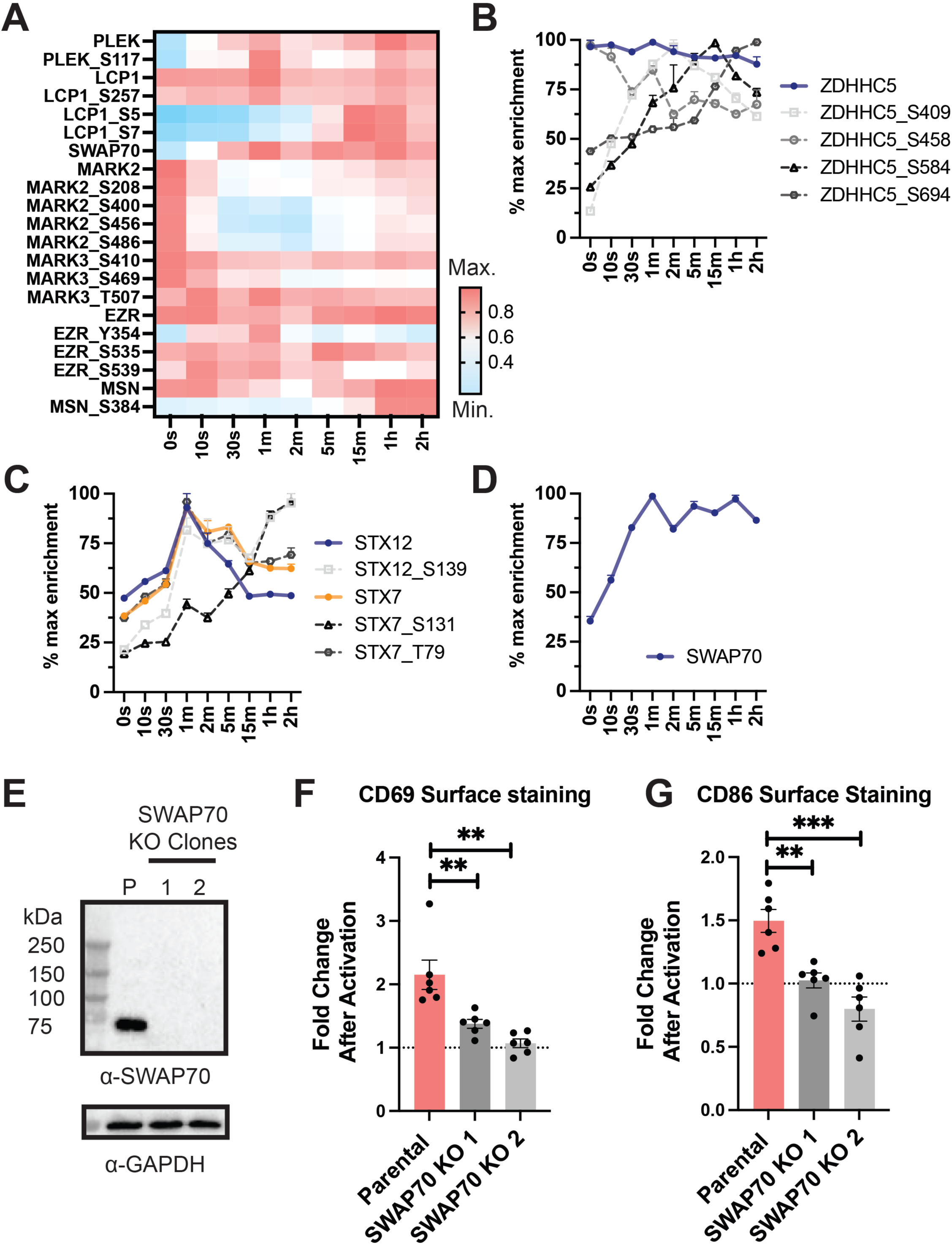
CD19-APEX2 as a resource to uncover new B cell biology. **(A)** Enrichment pattern heatmap of proteins involved in cytoskeletal regulation and organization. **(B)** Enrichment pattern heatmap of the palmitoyltransferase ZDHHC5 and several phosphorylation sites. Error bars represent mean ± SEM of two biological replicates. **(C)** Enrichment pattern heatmap of syntaxins (STX). Error bars represent mean ± SEM of two biological replicates. **(D)** Enrichment profile of SWAP70. Error bars represent mean ± SEM of two biological replicates. **(E)** Western blot of SWAP70 knockout Raji cells. “P” denotes parental Raji cells. **(F)** Fold change in CD69 surface staining after activation with anti-IgM F(ab’)2 in parental Raji cells and SWAP70 knockout clones. **(G)** Fold change in CD86 surface staining after activation with anti-IgM F(ab’)2 in parental Raji cells and SWAP70 knockout clones. For panel F and G, error bars represent mean ± SEM of three independent experiments. Statistical analysis was performed in GraphPad Prism using an unpaired t-test. *p ≤0.05; **p ≤ 0.01; ***p ≤ 0.001; ****p≤ 0.0001; ns, not significant. In all panels, phospho-marks are annotated with the protein name followed by “_amino acid position” of the phosho-mark.

In addition to proteins involved in cytoskeletal regulation, we identified dynamic changes in the abundance of phospho-marks on the palmitoyltransferase ZDHHC5 upon BCR stimulation. While there was no change in the relative amount of labeled ZDHHC5 throughout the time course, several phospho-marks exhibited some of the strongest changes in the dataset, increasing up to seven-fold upon activation **(Figure 4B).** Palmitoylation, a reversible modification to cysteine residues, regulates protein-lipid interactions to play an essential role in B cell activation. For example, a core component of the B cell co-receptor complex, the tetraspanin CD81, is palmitoylated upon B cell activation. This modification stabilizes the complex in cholesterol-rich regions of the membrane to enhance BCR signaling^60^. Notably, the enzymes that regulate the palmitoylation of specific proteins within immune cells are almost entirely unknown and uncovering the role of ZDHCC5 in B cell biology now allows for directly testing the hypothesis that it plays a functional role in B cell activation.

We also observe specific recruitment of Syntaxin 7 (STX7) and Syntaxin 12 (STX12) to the co-receptor complex at 1 minute following BCR activation **(Figure 4C)**. Syntaxins, a prototype family of SNARE proteins, are essential fusion mediators of both endocytic and exocytic trafficking events and are best understood for their roles in regulating neurotransmitter release. Syntaxin function has not been described in B cells. However, STX7 has been shown to mediate T cell receptor (TCR) recycling through endosomes, suggesting STX7 and STX12 may play a parallel role in B cells in BCR endocytosis^61^.

To highlight the ability of our system to identify physiologically relevant interactors, we examined if SWAP70, an actin-binding guanine nucleotide exchange factor (GEF), was necessary for B cell activation. SWAP70 has previously been linked to BCR signaling, specifically through its putative association with the BCR IgG isotype and as a component of a nuclear enzyme complex involved in immunoglobulin class switching^62–65^. SWAP70 has a low enrichment level before BCR crosslinking but is robustly recruited to CD19 within 30 seconds and remains in proximity of CD19 throughout the time course **(Figure 4D)**. These dynamics are consistent with a previous report showing cytoplasmic localization of SWAP70 in resting cells, plasma membrane localization in activated cells, and eventual nuclear localization several days after activation^63^. To investigate the relevance of SWAP70 in B cell activation, we created SWAP70 knockout Raji cells and then assayed for their ability to induce B cell activation markers in response to BCR stimulation **(Figure 4E).** SWAP70 knockout clones fail to increase CD69 and CD86 surface staining upon activation, indicating that SWAP70 is necessary for B cell activation in Raji cells **(Figure 4F and Figure 4G)**. The detection of dynamic changes in SWAP70 labeling by CD19-APEX further highlights the use of our proximity labeling approach to identify signaling components that play important yet poorly defined roles in B cell signaling.

### The glutamate transporter SLC1A1 mediates rapid glutamate uptake upon BCR stimulation

Our data set provides an unbiased resource for exploring and understanding the complex signaling networks underlying B cell activation, enabling the discovery of novel modulators. One particularly interesting and unexpected set of candidate regulators that emerged in our experiments is the solute carrier (SLC) family of transporters, which have no previously reported connections to B cell signaling. Specifically, we identified the glutamate transporter SLC1A1 (also known as EAAT3), a member of the excitatory amino acid transporter (EAAT) family, as a previously unidentified regulator of B cell activation.

In immune cells, solute carrier transporters (SLCs) transport various solutes (i.e. amino acids, lipids, ions) across the membrane to induce rapid metabolic reprogramming and nutrient uptake upon activation^66^. While SLCs are essential mediators of both normal and tumor cell function and are emerging as important immunotherapy targets to manipulate immunometabolism, the rational targeting of SLCs has been hindered by a poor understanding of which transporters mediate specific B cell functions^67^.

Although the metabolic requirements for B cell activation are relatively unknown^68, 69^, pharmacological manipulation of B cell metabolism could provide a powerful way to modulate B cell function and target aberrant proliferation, having implications for the treatment of B cell-driven autoimmune disease, B cell malignancies, and the development of vaccines ^68, 69^. Remarkably, 71 unique SLCs and phospho-marks were quantified in our proteomic dataset **(Figure 5A)**. Due to TMT multiplexing the data have almost no missing values, so we quantified the relative amount of specific site-phosphorylation in all experimental conditions in parallel. While the majority of SLCs did not change in abundance in the molecular neighborhood of CD19, several specific phosphorylation-sites showed inducible and sustained enrichment to the CD19 microenvironment. Phosphorylation on other sites over time, displayed depletion at late time points **(Figure 5A)**. Among the group of dynamic transporters, SLC1A1 stood out by becoming CD19-proximal within 1 minute after BCR stimulation and remaining proximal throughout the remaining time course **(Figure 5B).**

**Figure 5.**
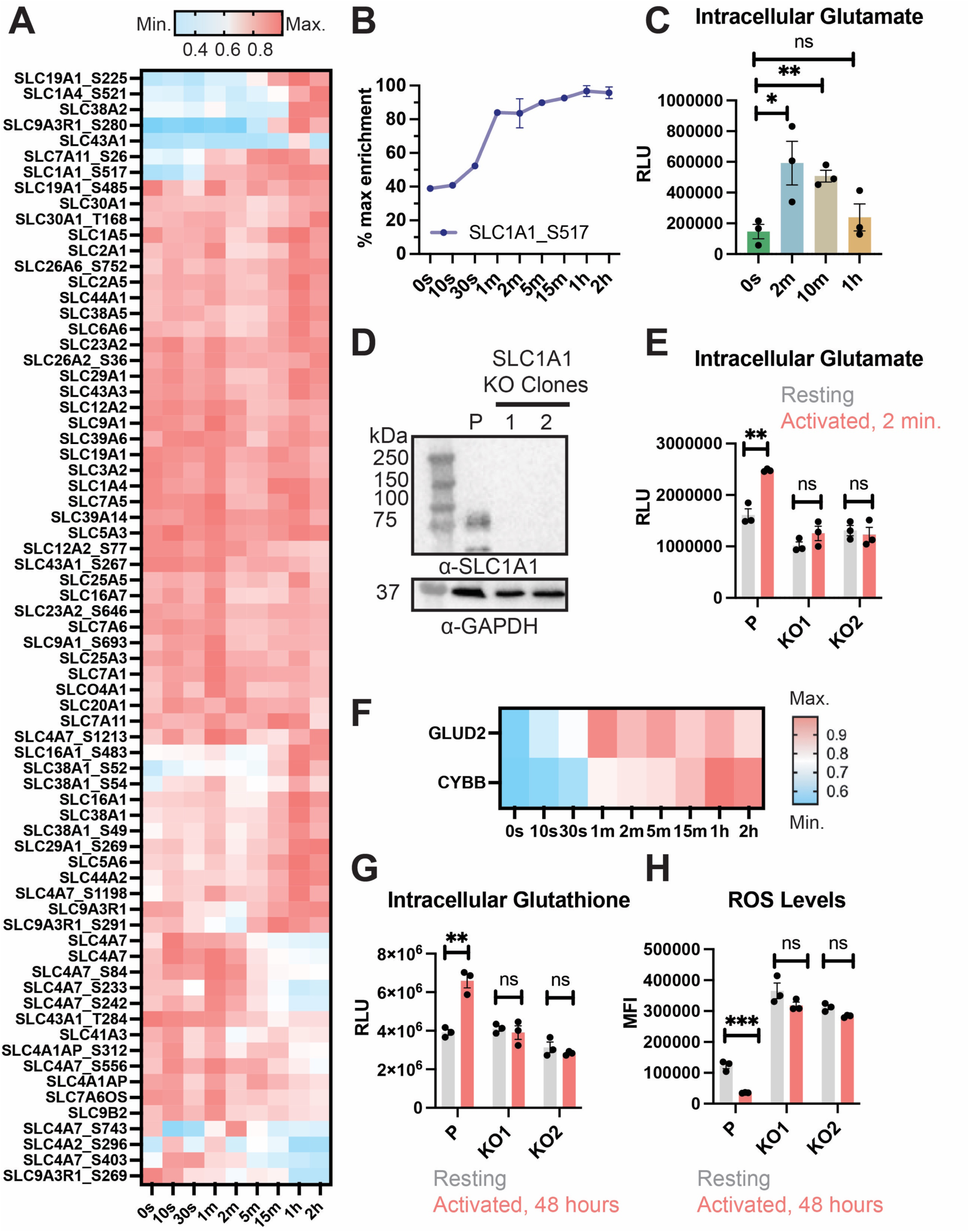
Recruitment and functional interrogation of the glutamate transporter SLC1A1. **(A)** Enrichment pattern heatmap of all members of the solute carrier (SLC) family identified in the CD19-APEX time course. Each protein’s enrichment level across different time points is normalized to its maximum enrichment signal. **(B)** Enrichment profile of the glutamate transporter SLC1A1. Error bars represent mean ± SEM of two biological replicates. **(C)** Glutamate uptake kinetics in parental Raji cells upon activation. **(D)** Western blot of SLC1A1 knockout Raji cells. “P” denotes parental Raji cells. **(E)** Glutamate uptake in parental and SLC1A1 knockout Raji cells upon activation for 2 minutes. **(F)** Enrichment pattern heatmap of NADPH oxidase 2 (CYBB) and glutamate dehydrogenase 2 GLUD2. Each protein’s enrichment level across different time points is normalized to its maximum enrichment signal. **(G)** Glutathione levels in parental and SLC1A1 knockout Raji cells in resting and activated cells, measured 48 hours after activation. **(H)** ROS levels in parental and SLC1A1 knockout Raji cells in resting and activated cells, measured 48 hours after activation. For Panel C, E, G, and H, error bars represent mean ± SEM of three independent experiments. Statistical analysis was performed in GraphPad Prism using an unpaired t-test. *p ≤0.05; **p ≤ 0.01; ***p ≤ 0.001; ****p≤ 0.0001; ns, not significant. In all panels, phospho-marks are annotated with the protein name followed by “_amino acid position” of the phosho-mark.

While SLC1A1 has well-described roles in maintaining glutamate levels within the brain^70^ and glutamate has some previously reported roles in T cell activation^71, 72^, the strong enrichment of SLC1A1 was unexpected because a role for glutamate uptake in the B cell activation response has not been reported. Using a luminescence-based assay to monitor intracellular glutamate levels, we showed that Raji cells rapidly take up glutamate upon activation, with levels increasing over three-fold from basal levels at 2 minutes and 10 minutes after BCR stimulation **(Figure 5C)**. To investigate whether this uptake depended on the presence of SLC1A1, we created SLC1A1 knockout Raji cells and then assayed glutamate uptake after activation **(Figure 5D)**. SLC1A1 knockout cells failed to increase intracellular glutamate, indicating that SLC1A1 is responsible for mediating rapid metabolic reprogramming immediately downstream of BCR stimulation **(Figure 5E)**.

Glutamate plays a critical role in central metabolism. It fuels the tricarboxylic acid (TCA) cycle, serves as a precursor for the synthesis of metabolites and other amino acids, and is the precursor for glutathione (γ-L-Glutamyl-L-cysteinyl glycine), an antioxidant tripeptide essential for scavenging reactive oxygen species (ROS) to maintain redox homeostasis in the cell. In line with these cellular functions, glutamate dehydrogenase (GLUD2), which catalyzes the reversible deamination of glutamate to alpha-ketoglutarate to feed into the TCA cycle, becomes highly enriched immediately after BCR crosslinking and remains enriched throughout the time course **(Figure 5F)**. Additionally, CYBB (also known as NOX2), an NADPH oxidase responsible for the generation of ROS, becomes strongly enriched beginning at 1 minute after BCR stimulation and remains CD19-proximal through the time course **(Figure 5F)**. Intracellular ROS is an important regulator of B cell activation^73, 74^. Upon activation, BCR stimulation triggers ROS production via NOX enzymes. The ROS reversibly oxidize phosphatases and limits their ability to dephosphorylate target kinases, which functions to amplify and prolong BCR signaling^75^.

We hypothesized that SLC1A1 may be involved in the regulation of redox homeostasis in B cells. Unlike other members of the EAAT family, SLC1A1 is unique in its ability to also transport cysteine, which is the rate-limiting intermediate for the biosynthesis of the main cellular antioxidant glutathione^76^. To determine if SLC1A1 is necessary for glutathione production in B cells, we assayed parental and SLC1A1 knockout Raji cells for their ability to synthesize glutathione upon activation. Parental cells showed a significant increase in glutathione levels 48 hours after BCR stimulation, while SLC1A1 knockout clones failed to synthesize glutathione **(Figure 5G)**. Parental Raji cells also showed a significant decrease in ROS amounts after long-term BCR stimulation, whereas SLC1A1 knockout cells have significantly increased amounts of ROS, which do not vary with B cell activation state **(Figure 5F)**. These data indicate that SLC1A1 mediates rapid metabolic reprogramming immediately upon BCR stimulation to maintain redox homeostasis during B cell activation.

## Discussion

Here, we used time-resolved peroxidase-catalyzed proximity labeling coupled with quantitative, multiplexed mass spectrometry to detail the sequential recruitment of signaling effectors to the endogenous B cell co-receptor complex in response to BCR stimulation. Whereas previous work investigating the dynamics of BCR signaling have analyzed temporal responses after five minutes or more with overexpressed proteins for co-immunoprecipitation or proximity labeling^36, 77, 78^, in this work we determined the kinetics of recruitment of previously known components of the BCR signaling pathway to CD19 tagged at its endogenous locus on a timescale from 10 seconds to 2 hours after BCR stimulation. Our experimental system revealed the kinetics of over 2,814 CD19-proximal proteins and 1,394 phospho-marks. While a substantial portion of the labeled proteins are bystanders that are in the same microenvironment but do not functionally associate with CD19, many others show striking changes in enrichment upon activation. Our unbiased and unprecedently extensive molecular map of the B cell activation pathway provides a unique resource both for understanding the complex signaling networks underlying B cell activation and for identifying potential targets for modulating B cell function.

CD19 has been classically viewed as a membrane-anchored adaptor protein^44^, but our dataset highlights the highly dynamic nature of the proteins recruited to the CD19 microenvironment. Robust changes in proximity enrichment occur as early as 10 seconds after BCR stimulation across a remarkably broad range of proteins, including kinases, second messenger generators, other co-receptors, and actin-binding proteins. Prior studies tracked BCR interactions beginning at 5 minutes after stimulation using co-immunoprecipitation followed by subsequent identification of interacting proteins with mass spectrometry^78^ or selective proteomic proximity labeling using tyramide (SPPLAT)^77^. More recently, an over-expressed APEX2 fusion was targeted to lipid rafts to monitor B cell signaling at 5, 10, and 15 minutes after BCR stimulation^36^. Our dataset highlights that some of the most interesting and striking changes in the B cell co-receptor microenvironment occur within *seconds*, suggesting that the sparse time sampling used by these prior studies failed to capture many important events within the BCR signaling pathway. Importantly, our approach also included an enrichment step for phosphorylated proximal peptides, revealing extensive reorganization of phosphorylation signaling networks immediately upon BCR stimulation. A limitation of this approach, however, is that some phospho-peptides are inherently difficult to detect using mass spectrometry and the minuscule amounts of multiplexed phosphopeptides, micro-enriched from limited streptavidin pulldown material, make detection and quantification technically very challenging to begin with, so that phospho-quantification data is incomplete. For example, phosphorylation on Tyrosine 531 of CD19 was not quantified in our dataset, even though we could detect its rapid induction upon BCR stimulation by western blotting **(Figure S2E)**.

The discovery of an essential function for SLC1A1 in rapid metabolic reprogramming upon B cell activation highlights our dataset’s ability to reveal new mediators of BCR signaling, in addition to providing a quantitative tracking of previously known BCR signaling effectors. Although the metabolic requirements for B cell activation are almost entirely unknown, manipulation of B cell metabolism could provide a powerful way to modulate B cell function and target aberrant proliferation, having implications for the treatment of B cell-driven autoimmune disease, B cell malignancies, and the development of vaccines^68, 69^. For example, the development of an SLC1A1 targeting antibody could offer a means of controlling B cell proliferation and metabolism in malignant B cells.

Establishment of the CD19-APEX system highlights the utility of time-resolved proximity labeling in elucidating the signaling dynamics of proteins tagged with APEX2 at the endogenous locus and in a native cellular system. Robust B cell responses can be generated against highly diverse antigens, and our dataset reflects the response to an oligomeric antigen through a T cell-independent activation process, in which B cell receptor crosslinking directly activates B cells. The establishment of the CD19-APEX will also enable profiling the response to other stimuli, including membrane-bound and T cell-dependent antigens, enabling identification of both shared features of B cell activation and pathway-specific processes. Some B cell-derived diseases are driven by the response to a particular antigen class. For example, stimulation of the innate immune response by extracellular DNA is a driver of systemic lupus erythematosus^5^, and identification of unique pathway-specific processes offer exciting possibilities for future therapeutic development.

## Methods

### Resource Availability

#### Lead Contact

Further information and requests for resources and reagents should be directed to and will be fulfilled by the lead contact Andrew Kruse.

#### Materials Availability

All plasmids and cell lines used in this study are available from the lead contact.

#### Data and code availability

Mass spectrometry data have been deposited in the PRIDE database with accession number PXD040262.

### Key resources table

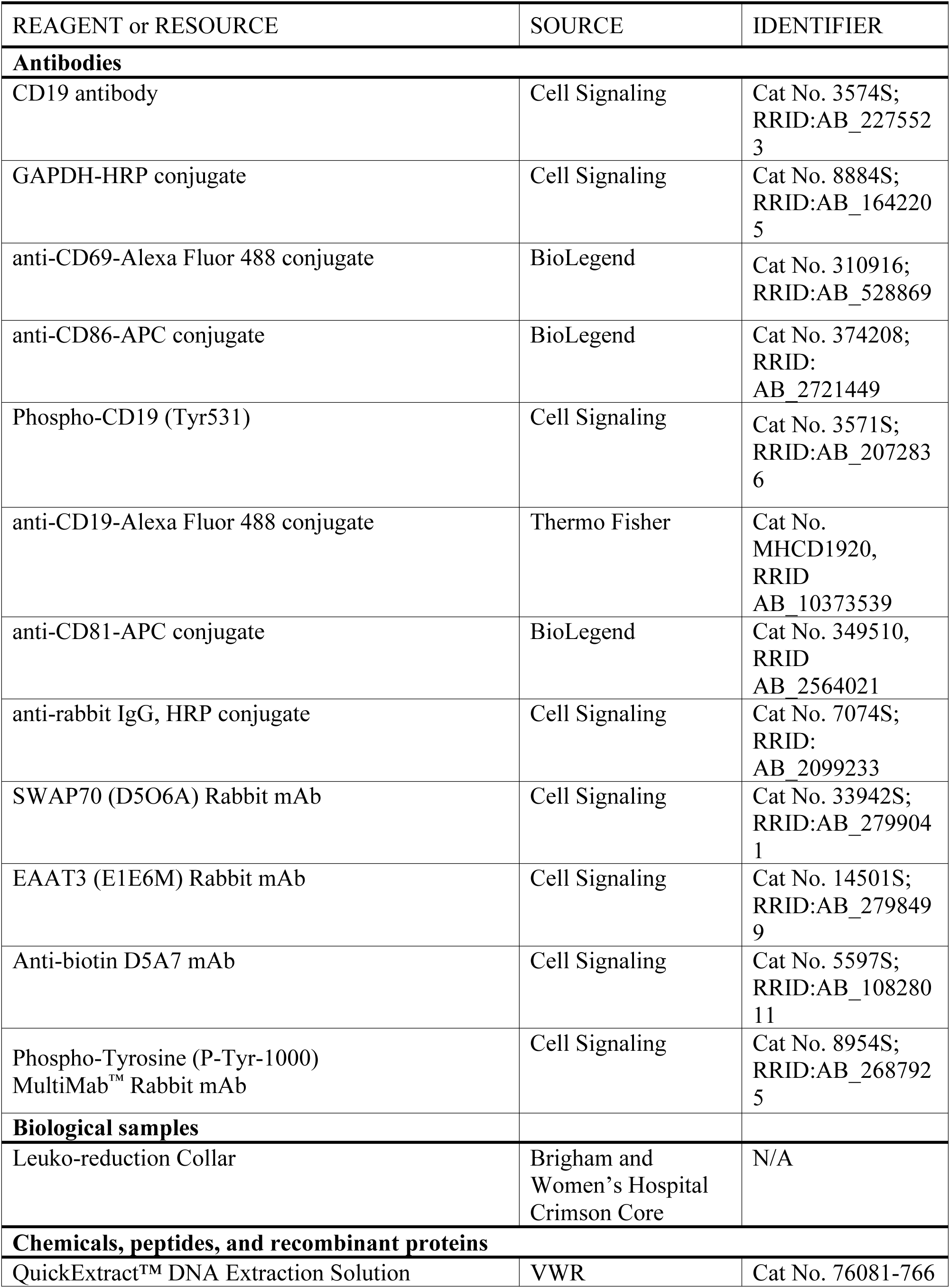
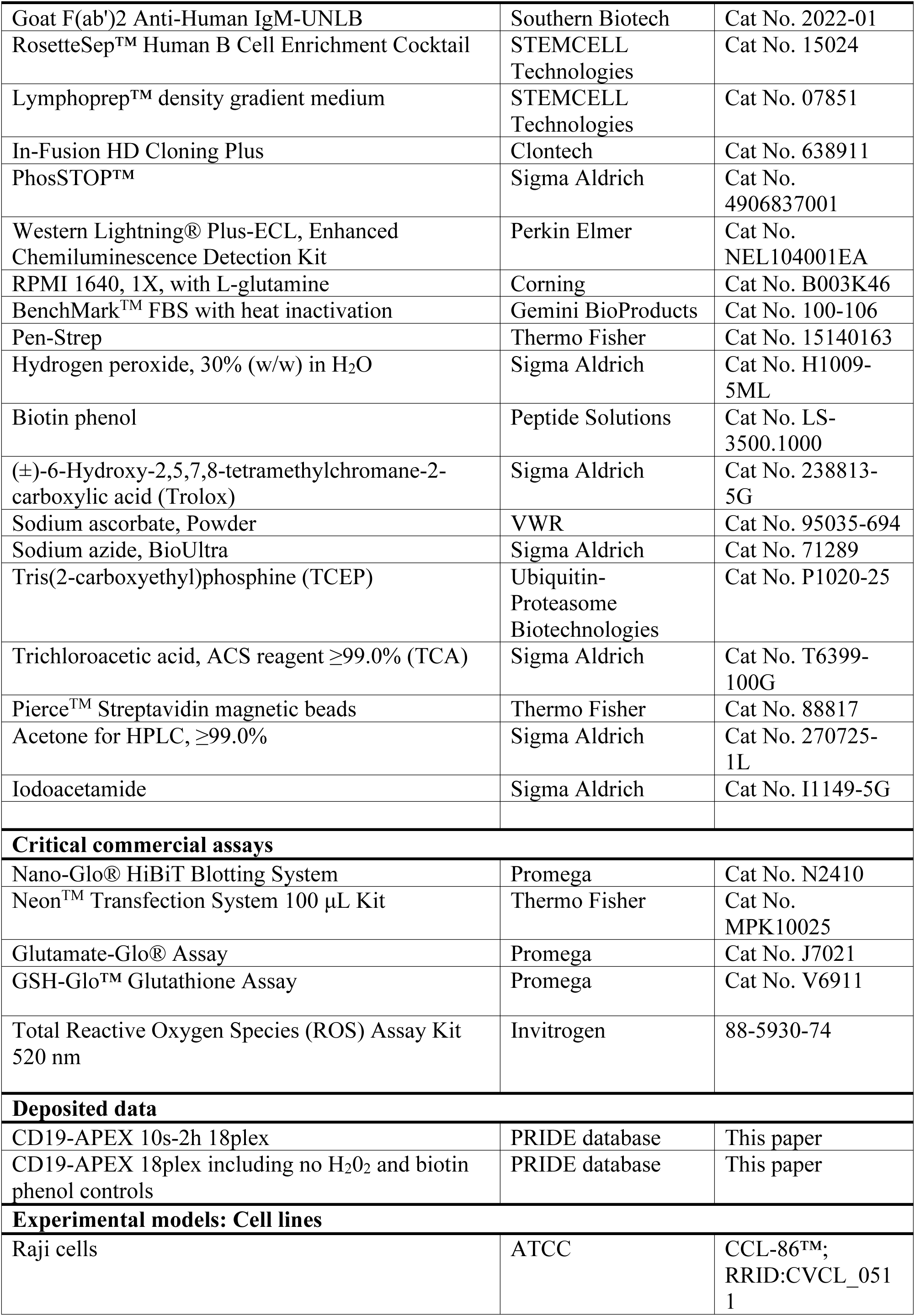
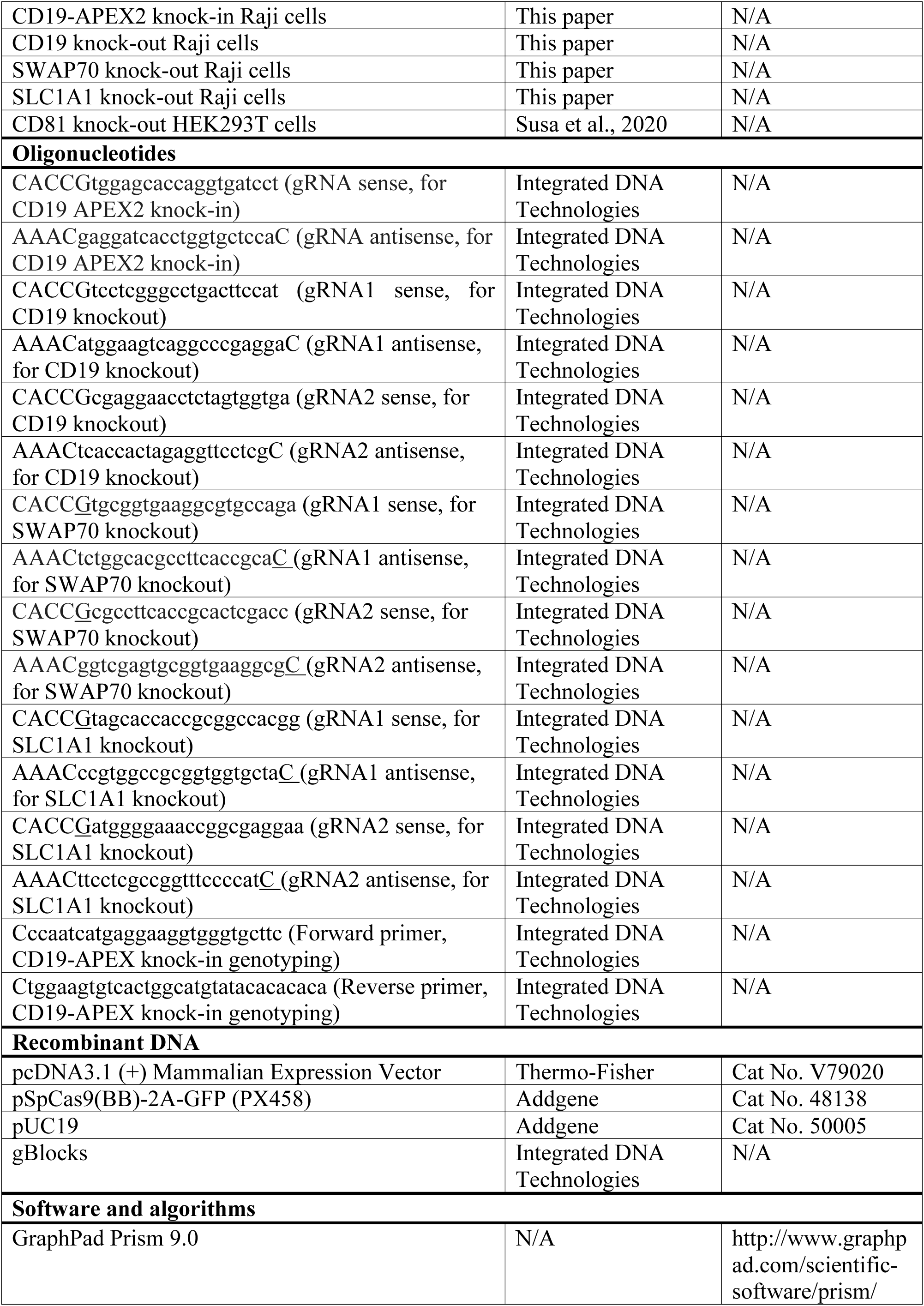
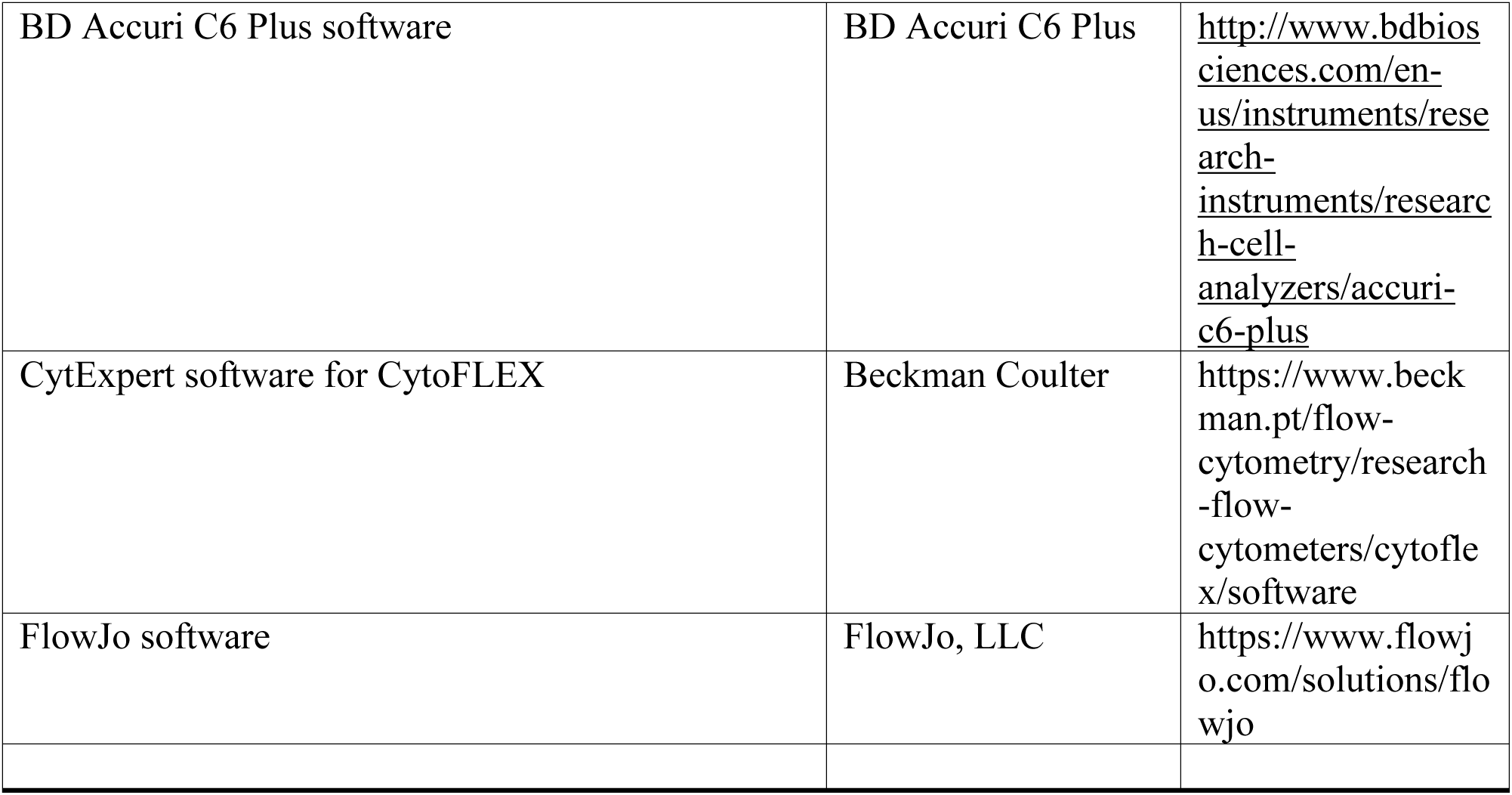

#### Engineering of CD19-APEX Raji Cells with CRISPR/Cas9

Raji cells were purchased from ATCC (CCL-86™) and confirmed to be negative for mycoplasma contamination using pooled primers and a previously reported PCR protocol ^79^. Raji cells were cultured at 37 °C and 5% CO_2_ in RPMI 1640 containing 10% fetal bovine serum (FBS; Gemini Bio-Products) and 1% Penicillin-Streptomycin (Pen-Strep; Thermo Fisher). To engineer a CD19-APEX knock-in cell line, Raji cells were co-transfected with the PX458-Cas9/GFP plasmid encoding a guide RNA (gRNA) targeting the sequence “TGGAGCACCAGGTGATCCTCAGG” at the CD19 C-terminus and a pUC19 plasmid encoding a CD19- GGSSGGSS-APEX2-HiBit repair template with ∼500 base pair homology arms. gRNAs were designed using the CRISPOR gRNA design tool (http://crispor.tefor.net/crispor.py), and the gRNA with the highest specificity scores was selected. Oligonucleotides sequences of gRNAs used for cloning into the PX458 plasmid are “CACCGtggagcaccaggtgatcct” (sense) and “AAACgaggatcacctggtgctccaC” (antisense).

The Neon transfection system (Thermo Fisher) was used for transfection. 10 million cells were spun down at 400 x g for 5 minutes at room temperature, washed once with 5 mL phosphate buffered saline (PBS), and resuspended in 200 µL Buffer R (Thermo Fisher). 30 µg of maxi-prepped DNA (Invitrogen) was added to the Buffer R-cell suspension, at a 3:1 molar ratio of repair template plasmid to Cas9 plasmid. Using a 100 µL tip, cells were then immediately electroporated at 1400 V for 20 ms with 2 pulses and added to 2 mL of pre-warmed RPMI 1640 containing 10% FBS and 1% Pen/Strep. In parallel, a mock transfection with the same parameters was performed without DNA as a negative control for setting cell sorting gates.

Two days after transfection, GFP^+^ single cells were sorted on a Sony SH800Z Sorter into 96 well plates containing 100 µL of media (RPMI containing 40% conditioned media, 40% fresh media, and 20% FBS). Two weeks later, single cell clones were first screened for insertion of an in-frame repair template using the Nano-Glo HiBit Lytic Detection System (Promega). HiBit-positive clones were then expanded and screened by anti-CD19 Western blot and genomic DNA PCR for genotyping.

For Western blots, 200 µL of cells at approximately 0.5 million cells/mL were spun down at 1000 x g for 3 minutes at 4°C. The supernatant was aspirated, and the cell pellet was resuspended in 50 µL non-reducing 2X SDS loading dye. Samples were separated by SDS-PAGE under non-reducing conditions, and the membrane was blocked with 5% w/v bovine serum albumin (BSA) in TBST (0.1% Tween-20 in Tris-buffered saline) at room temperature for 1 hour. The blocked membrane was cut and then incubated with shaking overnight at 4°C with either CD19 antibody #3574 (Cell Signaling; 1:1000 dilution) or GAPDH-HRP Conjugate (Cell Signaling, 1:1000 dilution) in TBST containing 1% BSA. The CD19 blot was then incubated with anti-rabbit IgG, HRP conjugate (Cell Signaling) diluted 1:5,000 in TBST containing 1% BSA for 1 hour at room temperature with shaking. Prior to chemiluminescent detection, blots were washed with TBST three times for 10 minutes each time. Western blots were developed with Western Lightning® Plus-ECL, Enhanced Chemiluminescence Detection Kit (PerkinElmer).

For clone genotyping, genomic DNA was extracted from individual clones using QuickExtract DNA solution (VWR) per manufacturer instructions. The CD19 gene was amplified by PCR using cccaatcatgaggaaggtgggtgcttc (forward primer) and ctggaagtgtcactggcatgtatacacacaca (reverse primer). Clones with the APEX2 insert showed a product at 1.3 kb, corresponding to an 850 bp increase compared to the parental cells, which showed a single band at approximately 0.5 kb. Bands were gel extracted and sequenced to confirm the presence of the APEX2 fusion. Off-target repair template integration in the genome was also assessed using the Nano-Glo® HiBiT Blotting System (Promega) per manufacturer instructions.

#### Flow cytometry assay for B cell activation

100 µL Raji cells were seeded at 0.5 million cells/mL in a 96 well plate approximately 18 hours before activation. Cells were either left unstimulated or 20 µg/mL of F(ab’)2 anti-human IgM (Southern Biotech) was added directly to the well. Approximately 48 hours later, cells were harvested at 300 x g for 3 minutes at 4°C, washed once with ice-cold PBS, stained with anti-CD69-Alexa Fluor 488 (Biolegend) and anti-CD86-APC (Biolegend) at 1:250 dilutions in 20mM HEPES buffer (pH 7.4), 150 mM NaCl, and 0.1% BSA for 30 minutes on ice, washed once with PBS, and then analyzed on a BD Accuri C6 flow cytometer or Beckman Coulter CytoFLEX.

#### Phospho-Tyrosine 531 CD19 assay

Parental Raji cells and CD19-APEX Raji cells were seeded at 0.5 million cells/mL, in a total volume of 1 mL for parental cells and 5 mL for knock-in cells, due to the lower abundance of CD19 in the knock-in cell line. Cells were returned to the incubator for 2 hours (37 °C and 5% CO_2_). Cells were then activated with 20 µg/mL of F(ab’)2 anti-human IgM (Southern Biotech) or left resting, and after 2 minutes cells were added to ice-cold PBS containing PhosSTOP™ (Sigma-Aldrich) to stop signaling. Samples were spun down at 1000 g for 2 minutes at 4°C, and the supernatant was aspirated. Cell pellets were resuspended in 40 µL ice-cold RIPA buffer containing PhosSTOP™ (Sigma-Aldrich), and then 15 µL of non-reducing SDS loading dye was added to each sample. 20 µL of sample was run on a 4-20% SDS PAGE gel at 110 V for 60 minutes. Proteins were then transferred to a nitrocellulose membrane at 25V for 1 hour at 4°C. Membranes were blocked with 5% w/v bovine serum albumin (BSA) in TBST (0.1% Tween-20 in Tris-buffered saline) at room temperature for 1 hour. The blocked membrane was then incubated with shaking overnight at 4°C with either CD19 antibody #3574 (Cell Signaling; 1:1000 dilution) or Phospho-CD19 (Tyr531) antibody (Cell Signaling, 1:1000 dilution) in TBST containing 1% BSA. The next morning, the blots were washed with TBST three times for 10 minutes each time. The blots were then incubated with anti-rabbit IgG, HRP conjugate (Cell Signaling) diluted 1:5,000 in TBST containing 1% BSA for 1 hour at room temperature with shaking. Prior to chemiluminescent detection, blots were washed with TBST three times for 10 minutes each time. Western blots were developed with Western Lightning® Plus-ECL, Enhanced Chemiluminescence Detection Kit (PerkinElmer).

#### CD19 Trafficking Assay

HEK293T cells (RRID CVCL_0063) were seeded at 100,000 cells/well in 24 well plates 12-18 hours prior to transfection. Cells were transfected using Lipofectamine 2000 with either with either 1.5 µg of empty pcDNA3.1(+) vector, 0.75 µg of CD19 DNA and 0.75 µg of empty pcDNA3.1(+) vector DNA (CD19 condition), or 0.75 µg of CD19 DNA and 0.75 µg of CD81 DNA (CD19+CD81 condition). 36-48 hours after transfection, cells were harvested in phosphate buffered saline (PBS) supplemented with 3 mM EDTA, transferred to a 96 well V-bottom plate, and then washed twice with PBS. Cells were then incubated on ice for 20 minutes with 2 µg/mL anti-CD19-Alexa Fluor 488 (Thermo Fisher) and anti-CD81-APC (BioLegend, Catalog number 349510, RRID AB_2564021) in 20 mM HEPES buffer pH 7.4, containing 150 mM NaCl, and 0.1% BSA. Cells were washed two times with PBS and analyzed on a Beckman Coulter CytoFLEX.

#### Glutamate Uptake Assay

100 µL of Parental Raji cells or SLC1A1 knockout Raji cells were seeded at 0.6 million cells/mL in a 96 well plate 3 hours before activation. Cells were either left unstimulated or 20 µg/mL of F(ab’)2 anti-human IgM (Southern Biotech) was added directly to the well. At 2 minutes, 10 minutes, or 1 hour after activation, cells were harvested at 300 x g for 3 minutes at 4°C and then washed twice with ice-cold PBS. Cells were then resuspended in 25 µL ice-cold PBS, then 12.5 µL of Inactivation Solution (0.6N HCl) was added to each well to rapidly stop metabolism and destroy reduced NAD(P)H dinucleotides. Five minutes later, 12.5 µL of Neutralization Solution (1M Tris Base) was added to each well, and then 50 µL of Glutamate Detection Reagent (Promega, Glutamate-Glo Kit) was added to each well, and the plate was incubated for 60 minutes at room temperature. Luminescence was recorded with a 0.3 second integration time on a GloMax luminometer system (Promega).

#### Glutathione Assay

100 µL of Parental Raji cells or SLC1A1 knockout Raji cells were seeded at 0.5 million cells/mL in a 96 well plate the evening before activation. The next morning, cells were either left unstimulated or 20 µg/mL of F(ab’)2 anti-human IgM (Southern Biotech) was added directly to the well. 48 hours later, intracellular glutathione levels were measured with the GSH-Glo™ Glutathione Assay kit (Promega), following the manufacturer’s protocol. Luminescence was recorded with a 0.3 second integration time on a GloMax luminometer system (Promega).

#### ROS Assay

100 µL of Parental Raji cells or SLC1A1 knockout Raji cells were seeded at 0.5 million cells/mL in a 96 well plate 3 hours before activation. Samples were either left untreated or treated with 20 µg/mL F(ab’)2 anti-human IgM (Southern Biotech). 48 hours later, 0.2 µL of 500x ROS Assay Stain (Invitrogen) was added directly to each well, and then each well was mixed well. Samples were incubated for 60 minutes in a 37°C incubator with 5% CO_2_, and then samples were analyzed on a Beckman Coulter CytoFLEX flow cytometer.

#### Generation of Knockout cells with CRISPR/Cas9

To generate CD19, SWAP70, and SLC1A1 knockout Raji cell lines, the CRISPOR guide design tool (http://crispor.tefor.net/crispor.py) was used to select guide sequences targeting exon 1 of the *CD19, SWAP70, or SLC1A1* gene. For each gene, two gRNAs were co-transfected, with one targeting the forward strand and the other the reverse strand of the genomic locus. gRNAs were cloned into the PX458-Cas9/GFP plasmid, using the oligonucleotides listed in Table 4.2.

The Neon transfection system (Thermo Fisher) was used for transfection. 10 million cells were spun down at 400g for 5 minutes at room temperature, washed once with 5 mL PBS, and resuspended in 200 µL Buffer R (Thermo Fisher). 15 µg of the forward-targeting gRNA plasmid and 15 µg of the reverse-targeting gRNA plasmid were added to the Buffer R-cell suspension. Using a 100 µL tip, cells were then immediately electroporated at 1400V for 20 ms with 2 pulses and added to 2 mL of RPMI 1640 containing 10% FBS and 1% Pen-Strep. In parallel, a mock transfection with the same parameters was performed without DNA as a negative control for setting cell sorting gates.

Two days after transfection, GFP^+^ single cells were sorted on a Sony SH800Z Sorter into 96 well plates containing 100 µL of media (RPMI containing 40% conditioned media, 40% fresh media, and 20% FBS). Single clones were then expanded and validated for protein knockout by Western blot. For Western blots, 200 µL of cells at approximately 0.5 million cells/mL were spun down at 1000 x g for 3 minutes at 4°C. The supernatant was aspirated, and the cell pellet was resuspended in 50 µL non-reducing 2X SDS loading dye. Samples were separated by SDS-PAGE under non-reducing conditions, and the membrane was blocked with 5% w/v bovine serum albumin (BSA) in TBST (0.1% Tween-20 in Tris-buffered saline) at room temperature for 1 hour. The blocked membrane was cut and then incubated with shaking overnight at 4°C with either CD19 antibody #3574 (Cell Signaling; 1:1000 dilution), EAAT3 antibody (Cell Signaling, 1:1000 dilution), SWAP70 antibody (Cell Signaling, 1:1000 dilution) or GAPDH-HRP Conjugate (Cell Signaling, 1:1000 dilution) in TBST containing 1% BSA. Blots were then incubated with anti-rabbit IgG, HRP conjugate (Cell Signaling) diluted 1:5,000 in TBST containing 1% BSA for 1 hour at room temperature with shaking. Prior to chemiluminescent detection, blots were washed with TBST three times for 10 minutes each time. Western blots were developed with Western Lightning® Plus-ECL, Enhanced Chemiluminescence Detection Kit (PerkinElmer).

#### Flow cytometry of CD19 surface expression

100 µL Raji cells or primary human B cells were seeded at 0.5 million cells/mL in a 96 well plate. Approximately 24 hours later, cells were harvested at 300 g for 3 minutes at 4°C, washed once with ice-cold PBS, stained with anti-CD19-Alexa Fluor 488 (Thermo Fisher) at 1:250 dilution in 20mM HEPES buffer (pH 7.4), 150 mM NaCl, and 0.1% BSA for 30 minutes on ice, washed once with PBS, and then analyzed on a Beckman Coulter CytoFLEX instrument.

#### Proximity labeling experiments

For each time point, CD19-APEX Raji cells were seeded at 1 million cells per mL in 10 mL of RPMI 1640 containing 10% FBS and 1% Pen-Strep in a T-25 flask, then returned to the incubator for two hours. Two hours before labeling, biotin phenol (Peptide Solutions) was added to the media at a final concentration of 2 mM. At various time points after addition of 20 µg/mL of F(ab’)2 anti-human IgM (Southern Biotech), cells were then labeled by addition of hydrogen peroxide (H_2_O_2_; Sigma Aldrich) at a final concentration of 0.25 mM. Immediately before each experiment, a new bottle of 30% (v/v) H_2_O_2_ stock solution was freshly diluted with PBS to make a 1 M working stock. After 1 minute of labeling with H_2_O_2_, the reaction was quenched by addition of 100 µL 1 M sodium ascorbate, 100 µL 1 M sodium azide, and 100 µL 0.5 M Trolox directly to the media. The cells were then immediately transferred to a falcon tube containing 10 mL of ice-cold Quench Buffer (PBS containing 10 mM sodium ascorbate, 5 mM Trolox, and 10 mM sodium azide), spun at 1000g for 2 minutes at 4°C, washed again with 1 mL Quench Buffer, spun at 1000 g for 2 minutes at 4°C, then washed once with 1 mL of cold PBS. Cell pellets were then flash frozen in liquid nitrogen and stored at -80°C until streptavidin pulldown.

For all proximity labeling experiments, 20 µg/mL of F(ab’)2 anti-human IgM (Southern Biotech) was added to the cell medium and incubated for the indicated length of time, and in all cases, the duration of incubation with H_2_O_2_ was fixed to a 1-minute labeling interval. For example, for the 5-minute time point, anti-human IgM was incubated with the cells for 5 minutes, and then H_2_O_2_ was added to cells starting at 5 minutes and ending at 6 minutes after activation.

#### Streptavidin pull-down of biotinylated proteins

All buffers used for streptavidin pull-down experiments were prepared freshly and filtered through 0.22 µM filters. Frozen cells were lysed in 800 µL cell lysis buffer (8M urea, 100 mM sodium phosphate pH 8, 1% w/v SDS, 100 mM ammonium bicarbonate, 10 mM TCEP), pipetted several times, and left on ice for 10 minutes. Samples were then loaded onto a QiaShredder (Qiagen) and spun at 21,130 x g for 2 minutes at room temperature. Proteins were then precipitated by addition of 800 µL of ice-cold 55% trichloroacetic acid (TCA) (Sigma Aldrich). Samples were left on ice for 15 minutes, and then spun at 17,000 g for 10 min at 4°C. All TCA was removed from the pellet with a gel loading tip, and then the pellet was washed with 1 mL of -20°C acetone, vortexed, and then incubated on ice for 10 minutes. Samples were centrifuged at 17,000 g for 10 min at 4°C, and then washed three more times with acetone. After the last acetone wash, all acetone was removed with a gel loading tip, and tubes were left open on ice for 15 min to remove residual acetone.

Dried cell pellets were resuspended in 1 mL cell lysis buffer and left on ice for 30 minutes, then heated to 37°C for 10 minutes to promote dissolving of the pellet. Re-suspended proteins were then centrifuged at 21,130 g for 10 minutes at room temperature, and the supernatant was transferred to a new microcentrifuge tube. Samples were then alkylated by addition of freshly prepared 400 mM iodoacetamide in 50 mM ammonium bicarbonate at a final concentration of 20 mM. The samples were then immediately vortexed and incubated in the dark for 25 minutes at room temperature. Dithiothreitol (DTT) was then added to the samples at a final concentration of 50 mM to quench the alkylation. 1 mL of water was then added to each sample to reach a final concentration of 4 M urea and 0.5% w/v SDS.

A 100 µL suspension per sample of streptavidin magnetic beads (Thermo Fisher Scientific) was washed twice with 4 M urea, 0.5% w/v SDS, 100 mM sodium phosphate pH 8 and was added to each sample. Tubes were rotated overnight at 4°C. Following streptavidin pulldown, magnetic beads were washed three times with 4 M urea, 0.5% w/v SDS, 100 mM sodium phosphate pH 8, three times with the same buffer without SDS, and three times in PBS. Beads were transferred to fresh tubes after the first wash in each set of buffers. After the final PBS wash, beads were stored at -80°C until mass spectrometry analysis.

#### On-bead digestion and tandem mass tag (TMT) labeling

Beads were transferred to clean tubes, and on-bead digestion was performed for 3 hours with the endoproteinase LysC and overnight with additional trypsin in 200mM EPPS buffer pH 8.5 with 2% acetonitrile at 37 °C under gentle agitation. Using a small aliquot of the digestion reaction, the missed trypsin cleavage rate was analyzed by mass spectrometry (MS), and for all reactions was less than 10%.

Digested peptides were then labeled with TMT 16plex (TMTpro, ThermoFisher cat# A44520) or TMT 18plex (TMTpro, ThermoFisher cat# A52045) reagents in enzyme digestion reactions (200mM EPPS pH 8.5) after addition of 30% acetonitrile (v/v). TMT labeling efficiency was determined by injecting 1% of stage tipped^80^, mixed labeling reactions. For all samples, labeling efficiency was over >95% as measured by N-terminal TMTpro conjugated peptides from MS data dynamically searched for TMTpro (+304.207 Da) peptide N-terminus modification and respective static modifications on lysine residues. TMT labeling reactions were then quenched by addition of 0.5% hydroxylamine for 15 minutes followed by acidification with formic acid, and then acetonitrile was evaporated by vacuum speed centrifugation (“speed-vac”) to near dryness. Next, peptides were resuspended in 1% formic acid and 0.1% Trifluoroacetic acid (TFA) and desalted and purified by reversed phase chromatography via C_18_ Sep-Pak cartridges (Waters #WAT054945) by gravity flow. Prior to the loading of TMT labeled peptides, C_18_ columns were washed with methanol, acetonitrile, then 1% formic acid with 0.1% TFA. Loaded peptides were washed with 3 column volumes of 1% formic acid and eluted with 95% acetonitrile with 1% formic acid. Peptides were dried to near completion and then subjected to subsequent micro-phospho enrichment.

#### Micro phospho-peptide enrichment

Dried peptides were re-suspended in 50 µL Fe-NTA binding buffer (0.1% TFA, 80% acetonitrile; included in High-Select^TM^ Fe-NTA Phosphopeptide Enrichment Kit, Thermo Scientific #A32992).

To create a micro Fe-NTA enrichment column, a small C18 matrix plug was inserted into a pipette tip after punching out a small amount of C18 matrix from a C18 Empore^TM^ solid phase extraction disk (Sigma-Aldrich #66883-U) with a 16 gauge blunt ended syringe needle, creating a one disk STAGE-tip as widely used in mass spectrometry ^80^. The tip was washed and conditioned by passing 100 µL of neat acetonitrile through, followed by addition of Fe-NTA matrix. For this, 20 µL of re-suspended Fe-NTA bead slurry from a High-Select^TM^ column was added into the STAGE-tip by pipetting with a cut-off pipet tip, resulting in transfer of approximately 10 µL of Fe-NTA beads. Bead storage buffer was passed manually and gently through the micro-enrichment tip by air pressure with a cut-off pipettor tip, preventing the beads from drying. Beads were washed by passing 70 µL of Fe-NTA binding buffer through the micro phospho-enrichment tip twice and re-suspended peptides in Fe-NTA binding buffer were added to Fe-NTA beads with a fine pipette tip minimizing contact with walls of the enrichment tip. Residual air bubbles between peptide solution and Fe-NTA beads were removed by gentle pipetting with a fine long-reach tip, also re-suspending Fe-NTA beads in the peptide solution. Fe-NTA beads were resuspended in the micro enrichment column every ten minutes for a total binding time of 30 minutes. After 30 minutes of binding, peptide solution was passed through the micro enrichment tip into a clean collection tube and collected together with 3 washes with 70 µL Fe-NTA binding buffer, a single wash with 70 µL ultrapure water and 50 µL of neat acetonitrile. The combined flowthrough and washes were dried down in a speed vac to completion for subsequent bench-top fractionation of non-phospho peptides by alkaline reversed phase chromatography (Pierce^TM^ High pH Reversed Phase Peptide Fractionation Kit, Thermo Scientific #84868) with a modified elution scheme as described in^81^ and MS analysis.

After binding to Fe-NTA beads, peptides were eluted with 25 µL of Fe-NTA elution buffer (ammonia solution in ultrapure water, pH 11.3; elution buffer provided with High-Select^TM^ kit) two times followed by elution with 50 µL neat acetonitrile. Beads turn dark brown after elution. Ammonia and acetonitrile eluates were collected directly in a single glass insert for MS autosampler vials, dried down to completion in a speed vac and dried phospho-peptides were re-suspended in 6 µL 1% formic acid for subsequent uHPLC injection and MS analysis.

#### Mass spectrometry methods

Data were collected on an Orbitrap Fusion Lumos instrument (Thermo Fisher Scientific) with a Multi Notch MS^3^ method ^82^ using a scan sequence of MS1 orbitrap scans (resolution 120,000; mass range 400-2000 Th), MS2 scans after collision-induced dissociation (CID, CE=35) in the ion trap with varying injection times. For phospho-peptide analysis, a multi-stage activation method was used with a neutral loss of 97.9763Da. Quantitative information was derived from subsequent MS3 scans (orbitrap, resolution 50,000 at 200 Th) after high-energy collision induced dissociation (HCD). Peptides were ionized by electrospray after HPLC separation with a MS-coupled Proxeon EASY-nLC 1200 liquid chromatography system (Thermo Fisher Scientific) using 75 µm inner diameter nanocapillaries with PicoTip emitters (NewObjective # PF360-75-10-CE-5) packed with C18 resin (2.6 µm, 150Å, Thermo Fisher Scientific). Fractionated peptides were separated over 3- and 4-hour gradients with increasing concentrations of acetonitrile from 0% to 95% in 0.125% formic acid. For phospho-peptide analysis, peptides were resuspended in 1% formic acid without acetonitrile prior to nano LC injection and separated over 120-minute and 60-minute gradients with 50% injected each as low amounts of material did not permit efficient fractionation. Injection times for phospho-peptides varied with up to 600ms for MS2 and 1000ms for MS3. A modified ANOVA score (“ModScore”) as described previously ^83^ was used to calculate localization confidence in phosphorylated residue position in the quantified phosphopeptides. MaxModscores of >13 correspond to confident localization of the phosphorylated residue position.

#### Western blotting analysis of biotinylated proteins

Samples of cell lysates were analyzed by Western blotting for biotinylation. After SDS-PAGE under non-reducing conditions, proteins were transferred to a nitrocellulose membrane and stained with Ponceau S (Sigma Aldrich) to monitor protein loading. Membranes were blocked in 5% BSA (w/v) in TBST for 1 hour at room temperature. Blocked membranes were then incubated with anti-biotin D5A7 antibody (Cell signaling) diluted 1:2,000 in TBST containing 1% BSA (w/v) overnight at 4°C. The membrane was then washed once with TBST for 5 minutes at room temperature, and then incubated with anti-rabbit IgG, HRP conjugate (Cell Signaling) diluted 1:5,000 in TBST containing 1% BSA for 1 hour at room temperature with shaking. Prior to chemiluminescent detection, blots were washed with TBST three times for 10 minutes each time. Western blots were developed with Western Lightning® Plus-ECL, Enhanced Chemiluminescence Detection Kit (PerkinElmer).

#### Purification of primary human B cells

A leuko-reduction collar was obtained from the Brigham and Women’s Hospital Crimson Core with patient information deidentified. All methods were carried out in accordance with relevant guidelines and regulations. All experimental protocols were reviewed and approved as exempt by the Harvard Faculty of Medicine Institutional Review Board. Primary human B cells were isolated from fresh leuko-reduction collar blood by using 750 µL of RosetteSep™ Human B Cell Enrichment Cocktail (Stemcell Technologies) for 10 mL of collar blood. The RosetteSep™ cocktail was incubated with collar blood for 20 minutes at room temperature, and then 10 mL of PBS supplemented with 2% FBS was added to the collar blood and mixed gently. The diluted collar blood was then layered on top of 10 mL Lymphoprep™ density gradient medium (Stemcell Technologies). After centrifugation at 1200 g for 20 minutes, the mononuclear cell layer was harvested and washed twice with PBS supplemented with 2% FBS. Purified cells were then frozen in 80% FBS and 20% DMSO at 10^7^ cells/mL until use. Upon thawing, cells were resuspended in warm RPMI 1640 supplemented with 10% FBS and 1% Pen-Strep at 10^6^ cells/mL.

## Acknowledgments

We thank members of Blacklow and Kruse labs for helpful discussions, and Jason Cyster for a critical reading of the manuscript. Financial support for this work was provided by NIH grants R01AI172846 (to S.C.B. and A.C.K), R35CA220340 (S.C.B.), F31HL147459 (K.J.S.), and a UCSF Sandler Fellowship (K.J.S.).

## Author Contributions

K.J.S., S.C.B., and A.C.K. conceived the project. K.J.S. performed time course experiments and cell-based assays. K.J.S. generated and validated CRISPR-edited cell lines with assistance from C.M.S. G.A.B., R.J.E., and M.K., processed and analyzed mass spectrometry data. K.J.S, S.C.B., and A.C.K. wrote the manuscript with input from all authors.

