## Supplementary_information for "A Spatiotemporal Map of Co-Receptor Signaling Networks Underlying B Cell Activation"

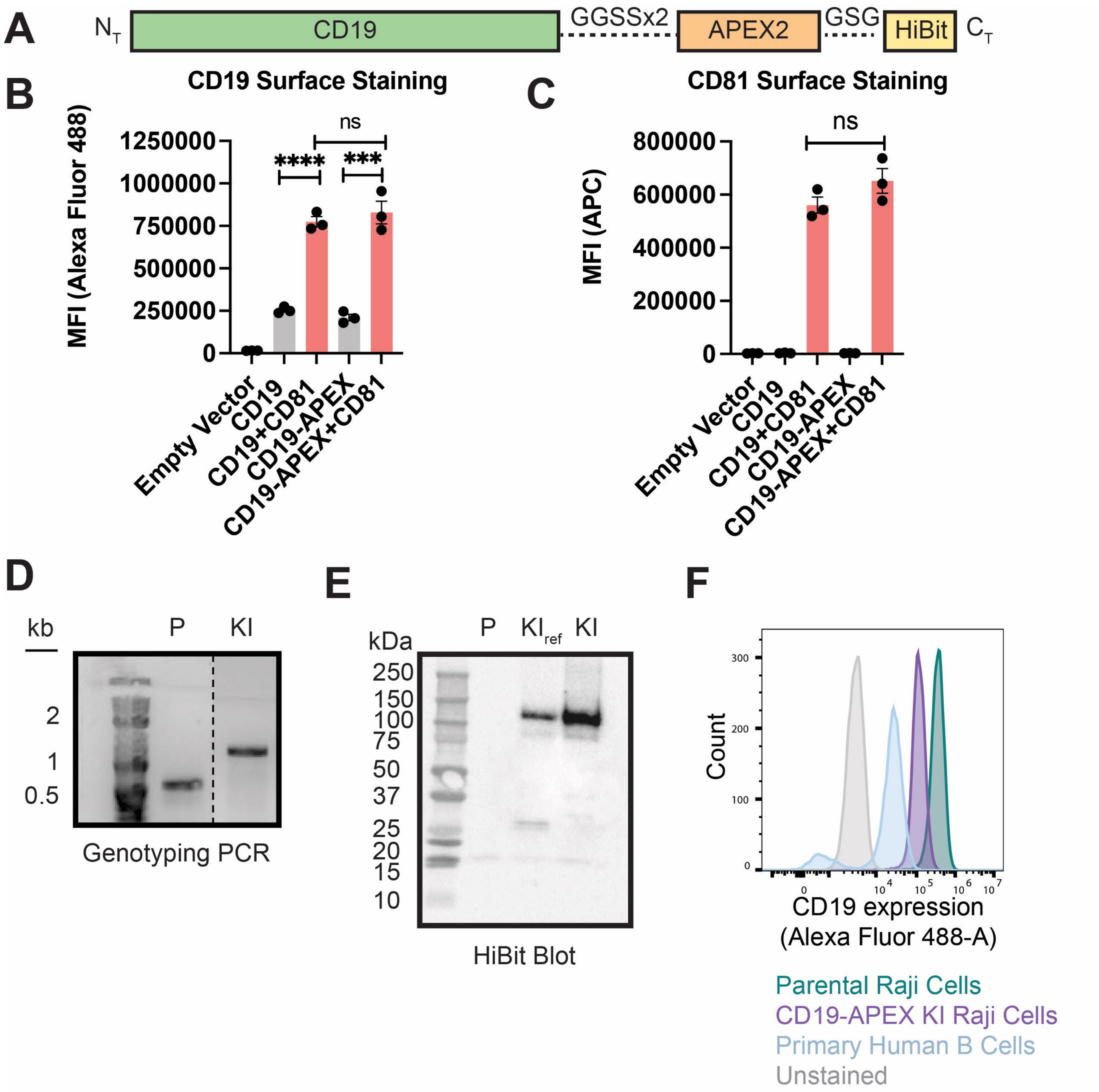


**Figure S1: Engineering and Validation of CRISPR-edited CD19-APEX2 Raji cells. Related to Figure 1. (A)** Schematic of CD19-APEX2 fusion construct. **(B)** CD19 export assay in CD81 knockout HEK293T cells. **(C)** Surface expression of CD81 constructs used in the export assay shown in Panel B. **(D)** Genotyping PCR of homozygous CD19-APEX clone used for all labeling experiments. "P" denotes Parental Raji cells and "KI" denotes knock-in. **(E)** HiBit western blot to assess CD19-APEX2 integration specificity. The clone used in all experiments ("KI") shows a single band at the molecular weight of CD19, while an additional reference clone that was not used in any experiments, $KI_{ref}$, shows a minor band at 25 kDa, likely indicating the integration of APEX2 into an additional genomic locus. **(F)** CD19 surface staining of parental Raji cells, KI Raji cells, and primary B cells. Surface CD19 was detected by flow cytometry using an Alexa 488-coupled anti-CD19 antibody. For Panels B and C, error bars represent mean ± SEM of three independent experiments. Statistical analysis was performed in GraphPad Prism using an unpaired t-test. *p ≤0.05; **p ≤ 0.01; ***p ≤ 0.001; ****p≤ 0.0001; ns, not significant.

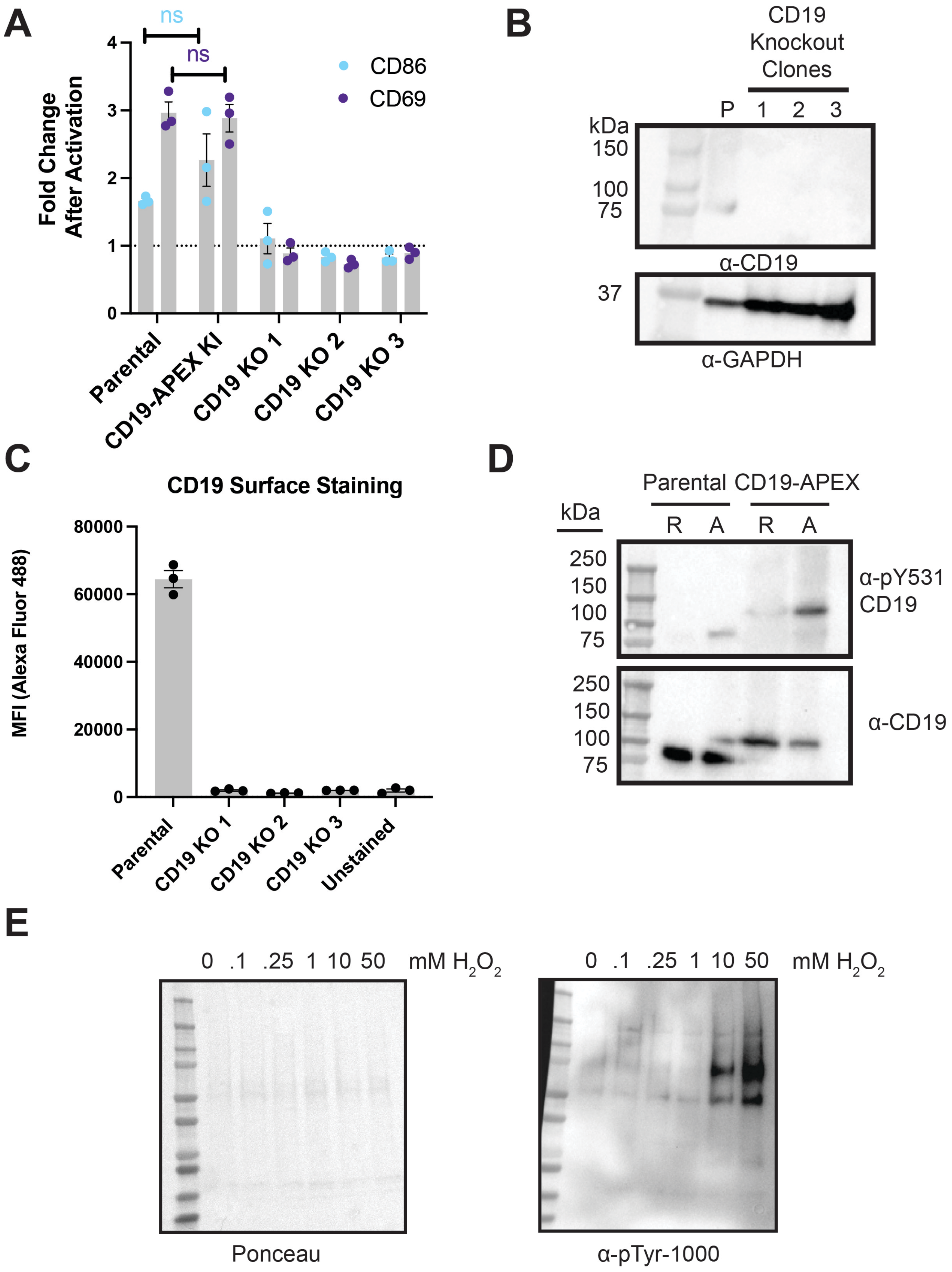
A
ns
ns
Fold Change After Activation
CD86
CD69
Parental
CD19-APEX KI
CD19 KO 1
CD19 KO 2
CD19 KO 3
B
CD19 Knockout Clones
kDa
P 1 2 3
150
100
75
α-CD19
37
α-GAPDH
C
CD19 Surface Staining
MFI (Alexa Fluor 488)
80000
60000
40000
20000
0
Parental
CD19 KO 1
CD19 KO 2
CD19 KO 3
Unstained
D
Parental CD19-APEX
kDa
R A R A
250
150
100
75
α-pY531 CD19
α-CD19
E
0 .1 .25 1 10 50 mM H2O2
Ponceau
α-pTyr-1000

**Figure S2: Validation of CD19-APEX2 Raji cell line in response to B cell activation. Related to Figure 1.** **(A)** Surface staining of the B cell activation markers, CD86 and CD69, in resting B cells and cells activated with anti-IgM F(ab')2. **(B)** Anti-CD19 western blot of the three CD19 knockout clones. "P" denotes parental Raji cells. **(C)** CD19 surface staining of three CD19 knockout clones, assessed by flow cytometry using an Alexa 488-coupled anti-CD19 antibody. **D)** Induction of phosphorylation on Tyrosine 531 of CD19 at 2 minutes after activation with anti-IgM F(ab')2. "R" denotes resting cells and "A" denotes activated cells. **(E)** Phospho-tyrosine Western Blot of Raji cells treated with 0, 0.1, 0.25, 1, 10, and 50 mM of $H_2O_2$. For Panels A and C, error bars represent mean ± SEM of three independent experiments. Statistical analysis was performed in GraphPad Prism using an unpaired t-test. *p ≤0.05; **p ≤ 0.01; ***p ≤ 0.001; ****p≤ 0.0001; ns, not significant.

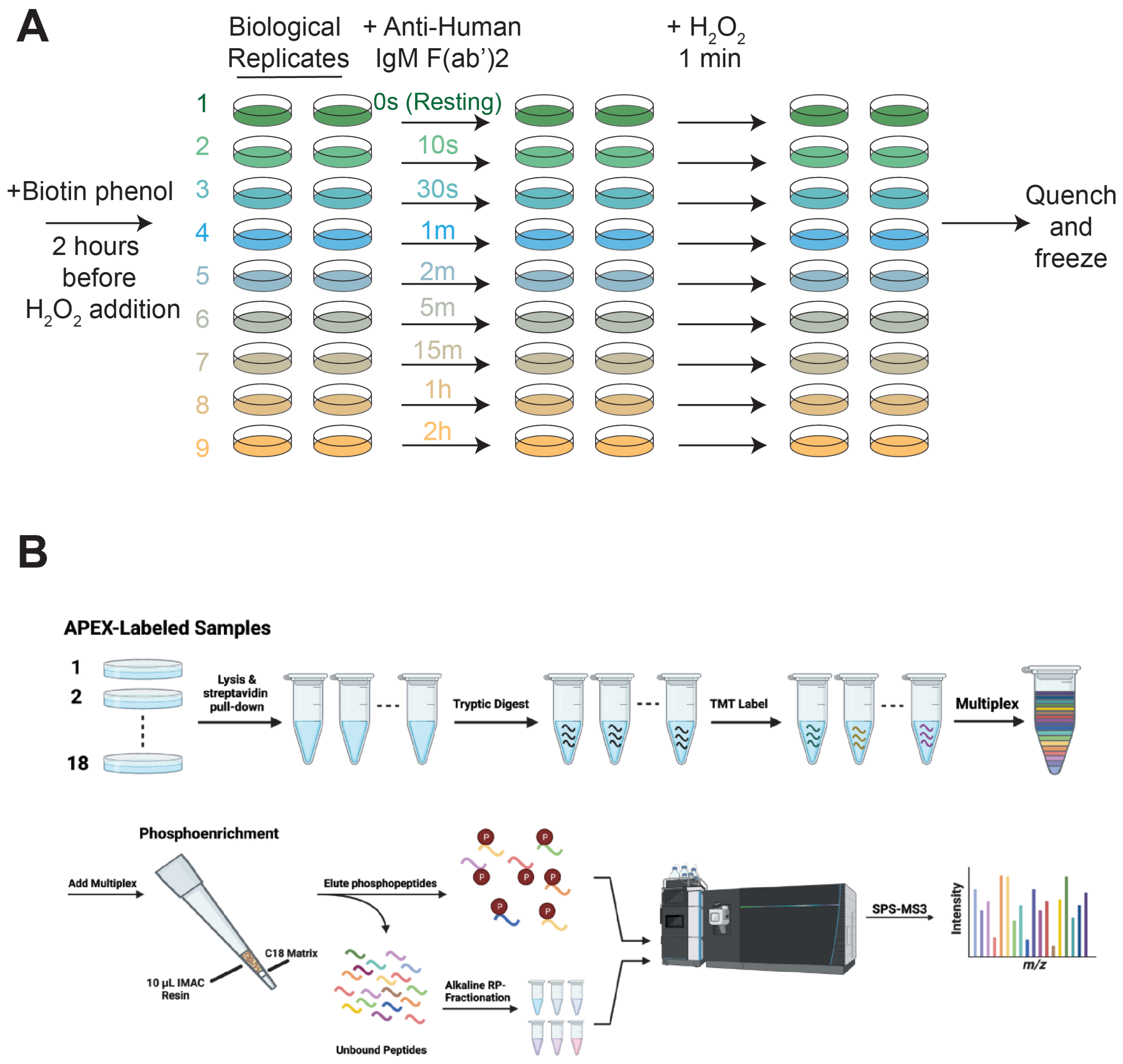


**Figure S3. Time course and mass spectrometry workflow. Related to Figure 2. (A)** Schematic of proximity labeling time course experiment. Cells were labeled with $H_2O_2$ for 1 minute at 10 seconds, 30 seconds, 1 minute, 2 minutes, 5 minutes, 15 minutes, 1 hour, or 2 hours after activation with anti-IgM (Fab)'2, or they were labeled with $H_2O_2$ without activation ("resting"). **(B)** Mass spectrometry workflow. Up to 18 samples are labeled with peroxide in the presence of biotin phenol at distinct time points after BCR stimulation. Biotinylated proteins are then purified under denaturing conditions with immobilized streptavidin, digested with trypsin, and then labeled with tandem mass tags (TMTs), enabling quantitative mass spectrometry identification of all samples in parallel. To identify and quantify phosphosites on biotinylated peptides, samples are also enriched using a Fe-NTA matrix.

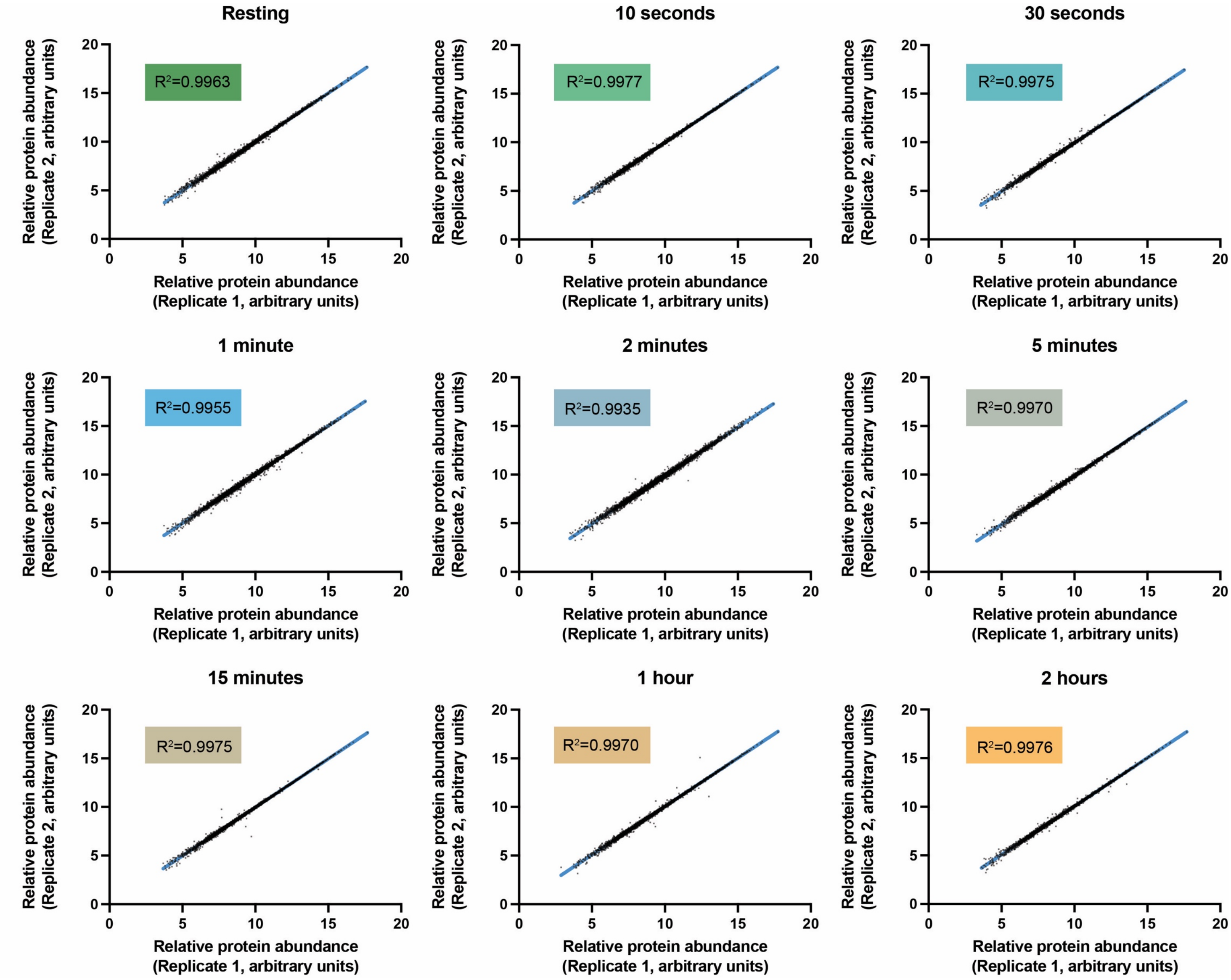


**Figure S4. Reproducibility of biological replicates. Related to Figure 2.** Representative linear regression analysis of protein and phosphomark abundance in biologically independent replicates labeled confirming high reproducibility in abundance measurements. Protein abundance is plotted on a log2 scale.

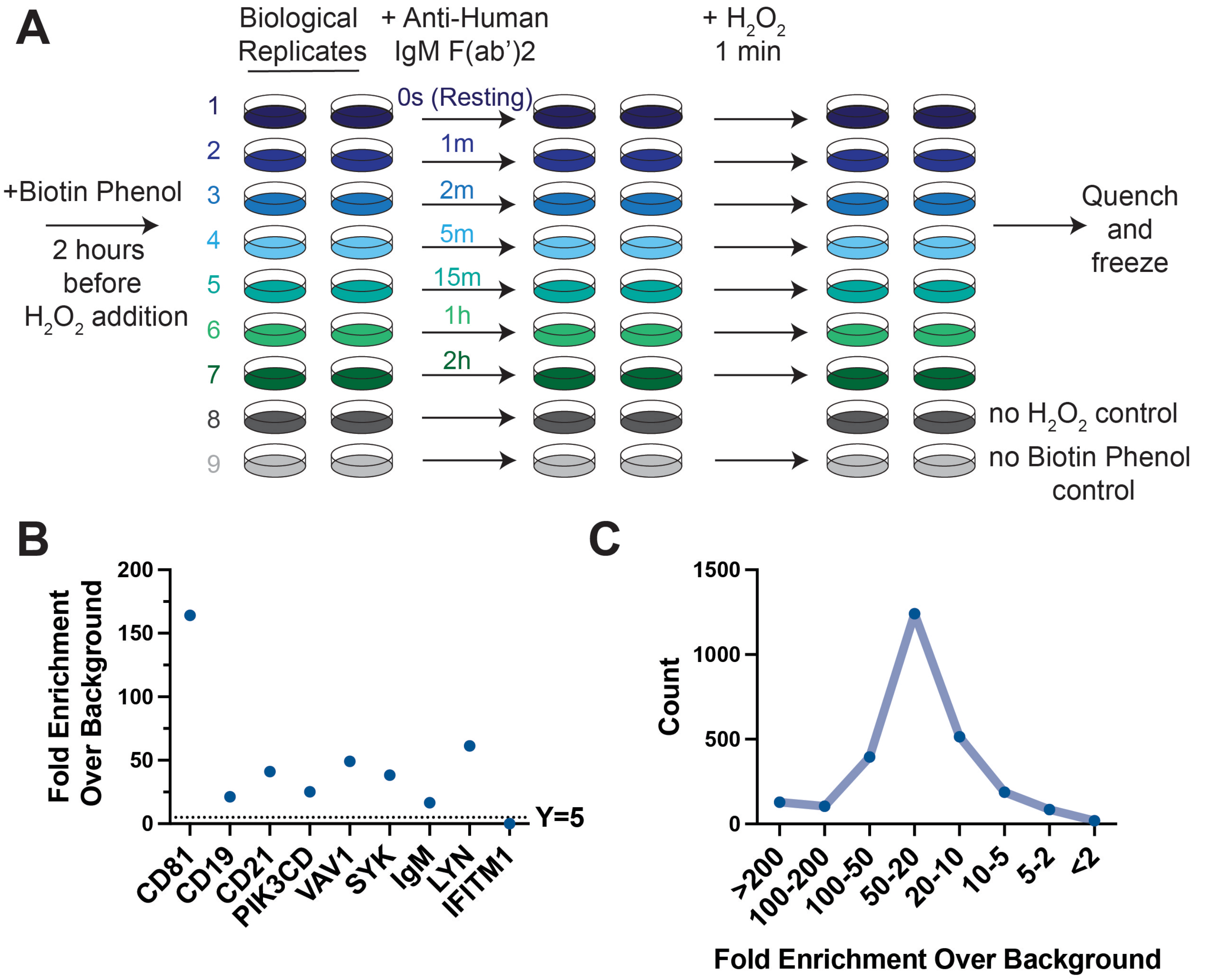


**Figure S5: Initial time course experiment of CD19-APEX2 cells stimulated with anti-IgM F(ab')2 with background labeling controls included. Related to Figure 2. (A)** Schematic of proximity labeling time course experiment. Cells were activated with anti-IgM F(ab')2 for 1 minute, 2 minutes, 5 minutes, 15 minutes, 1 hour, or 2 hours, or left untreated ("resting"). Controls omitting peroxide and biotin phenol treatment were also included to assess the extent of background labeling. **(B)** Fold enrichment of previously identified CD19 interactors in resting cells relative to background levels measured in cells without biotin phenol and peroxide treatment. CD19 interactors were identified from the STRING database. **(C)** Fold enrichment over background of all proteins identified in the time course.
